# Selective inflammation of the tumor microenvironment and invigorated T cell-mediated tumor control upon induced systemic inactivation of TREX1

**DOI:** 10.1101/2024.06.09.598086

**Authors:** Emilija Marinkovic, Minyi Chen, Nadja Schubert, Elif Dogan Dar, Janet Y. Leung, Jack Lohre, Jennifer M. Sahni, Christine Tun, Pavithra Rajeswaran, Tanja Mehlo-Jensen, Olivia Perng, C. Mark Hill, Pallavur Sivakumar, Michael Barnes, Rohit Malik, Rayk Behrendt, Axel Roers

**Affiliations:** Institute for Immunology, University Hospital Heidelberg, Heidelberg, Germany; DKMS Life Science Lab gGmbH, Dresden, Germany; Medical Systems Biology, Medical Faculty Carl Gustav Carus, TU Dresden, Dresden, Germany; Bristol Myers Squibb, Redwood City, CA, United States; Bristol Myers Squibb, Seattle, WA, United States; Institute for Clinical Chemistry and Clinical Pharmacology, University Hospital Bonn, Bonn, Germany

## Abstract

Therapeutic innate immune stimulation within the tumor microenvironment can potentiate endogenous antitumor T cell immunity. DNase 3’-repair exonuclease 1 (TREX1) is essential for cellular DNA disposal which prevents autoimmunity ensuing from cGAS/STING activation by endogenous DNA. Optimal strategies to therapeutically leverage cGAS/STING signalling for cancer therapy are highly sought after. TREX1-deficient tumor cells elicit enhanced protective immunity in syngeneic models. Here we show that induced inactivation of the *Trex1* gene in (non-malignant) host cells is well tolerated and yields improved type I IFN- and T cell-dependent control of established TREX1-competent tumors with selective immune cell infiltration of tumor, but not other tissues. Intra-tumoral T cell proliferation and numbers of effector and effector-like ‘exhausted’ cells massively increased, enabling complete rejection in synergy with checkpoint inhibition. We conclude that systemic TREX1 inhibition is a promising approach to boost anti-tumor immunity, that can overcome immune evasion by cancer cell-intrinsic cGAS/STING inactivation.

## Introduction

Amplification of endogenous T cell responses represents a central goal in cancer immunotherapy which has received intense attention in recent years due to the success of immune checkpoint inhibitor (ICI) therapies. These can achieve impressive long-term remission of various cancer entities, even including patients with end-stage disease (1). However, only a minority of patients benefit from ICI therapy, often those whose tumors exhibit significant dendritic cell (DC) and T cell infiltration prior to therapy. In contrast, sparse immune cell infiltration, defining immunologically ‘cold’ tumors, is a predictor of poor responsiveness to ICI therapy (1, 2). This limitation prompted a search for additional immunotherapeutic approaches synergizing with ICI therapies. As protective adaptive responses depend on innate signals, modulation of innate immunity is an important means to boost immune control of tumors (3, 4).

In this context, a focus of current interest is the cGAS/STING pathway that detects infection and cellular stress or damage to induce type I interferon (IFN) and pro-inflammatory cytokine responses (5, 6). Upon detection of double-stranded DNA of microbial or endogenous sources, the sensor cGAS catalyzes formation of 2’3’-cGAMP which is sensed by the ER-resident adapter STING. STING signals drive expression of type I IFNs as well as NF-κB-dependent inflammation (7), thereby stimulating adaptive immunity by enhancing DC maturation and antigen presentation, including cross-presentation and T cell co-stimulation. Accordingly, STING activation can efficiently adjuvant adaptive anti- microbial (8) and anti-tumor responses (5, 6).

Cancers often spontaneously mount tumor cell-intrinsic STING responses, as their genetic instability results in the release of different species of DNA into the cytosol. Genotoxic therapy enhances cytosolic DNA accumulation in tumor cells (9), and the cGAS/STING pathway has been shown to contribute to the immune-stimulatory effects of chemotherapy or irradiation (9, 10). Cancer cell-intrinsic cGAS activation and transfer of cGAMP to host immune cells of the TME can promote immune control of tumors (11, 12). Additionally, chromatin debris from dying cancer cells stimulates STING in host myeloid cells upon endocytosis, thereby promoting tumor control (13). Growth of syngeneic tumors is accelerated in hosts lacking STING (11, 14), and efficient tumor control by ICI has been shown to depend upon a functional STING pathway in host cells (14, 15).

Importantly, however, STING activation can also counteract anti-tumor immunity. For example, T cells express high levels of STING and are prone to STING agonist-induced cell death (16, 17), a mechanism that cancer cells can exploit for immune evasion by releasing cGAMP (18, 19). In poorly immunogenic tumors, STING was shown to suppress immune control via indoleamine 2,3-dioxygenase 1 activation (20) and myeloid-derived suppressor cell mobilization (21). The net effects of STING on immune responses depend on signal intensity, cell type and tissue context, and ideal approaches for therapeutic activation of this pathway, optimally leveraging its immune-adjuvanting potential, are currently an active field of research (5).

Both natural and synthetic STING agonists have been tested via intra-tumoral or systemic administration routes in various preclinical cancer models, overall demonstrating promising potential (reviewed in (5)). Intra-tumoral injections can be limited by tumor accessibility, local toxicity and the induction of T cell death by locally high concentrations of STING agonists, whereas systemic administration bears risks of acute inflammatory complications. Novel approaches to overcome these problems include, among others: orally deliverable non-nucleotide STING agonists, that were well-tolerated and effective in syngeneic cancer models (reviewed in (5)); innovative packaging of STING agonists in virus-like particles, exosomes or nanoparticle formulations (reviewed in (5)); and coupling of STING agonists to antibodies that target tumor-expressed surface structures (22). To date, clinical studies with STING agonists alone or in combination with ICI have shown moderate efficacy. This was partly attributed to STING activation in B cells, converting them into immunosuppressive IL-35^+^ regulatory B (Breg) cells (23).

An alternative strategy to activate STING is interference with a key counter-regulator of cGAS activation by endogenous DNA. Cytosolic DNA disposal critically depends on the ubiquitously expressed DNase 3’ repair exonuclease 1 (TREX1)(24, 25). Genetic defects of *TREX1* result in the accumulation of cytosolic DNA and chronic STING signaling that persistently stimulates adaptive immunity to cause the systemic lupus-like autoimmune disease, Aicardi-Goutières syndrome, a prototypic type I interferonopathy (26, 27). This autoinflammatory mechanism can be leveraged to stimulate T cell immunity. Murine studies demonstrated that TREX1-mediated cytosolic DNA disposal limits induction of systemic anti-tumor immunity upon high dose irradiation (28), and very recently three studies showed that loss of TREX1 in cancer cells can improve immune control of transplanted syngeneic mouse tumors, suggesting cancer cell-expressed TREX1 as a compelling therapeutic target (29–31).

As efficient inactivation of TREX1 in tumor cells can pose a challenge, and as cancers can inactivate the cGAS/STING axis or other key proteins of the type I IFN system to evade anti-tumor immunity (32–35), we studied effects of systemic interference with TREX1 activity in host cells. We show that induced inactivation of the *Trex1* gene in mice bearing established TREX1-proficient tumors results in selective inflammation of tumors, but not other tissues, and boosts CD8^+^ T cell-mediated tumor control. Combination of induced loss of TREX1 with ICI enabled complete tumor destruction. Our findings suggest that systemic pharmacologic inhibition of TREX1 activity is a promising strategy for boosting tumor immunotherapy that cancers cells cannot evade by silencing their cGAS/STING axis.

## Results

### Unaffected general health, but improved tumor control upon induced loss of TREX1 in host cells

To explore the potential of targeting TREX1 in non-malignant host cells to enhance tumor immune control as well as potential adverse effects, we made use of our *Trex1^FL/FL^R26^CreERT2^*mouse model (36) that allows for tamoxifen (TAM)-induced, systemic inactivation of the *Trex1* gene (‘TREX1 iKO’). Mice were treated with TAM via different routes, resulting in deletion of the *loxP*-flanked *Trex1* allele and subsequent loss of TREX1 protein in splenocytes (Fig. S1A,B). We chose intraperitoneal injection as the application route for all subsequent experiments. To study kinetics of the type I IFN response upon induced loss of TREX1, we collected spleens at different time points after a single injection of TAM and quantified ISG expression. *Ifi44* (Fig. 1A) and other ISG (not shown) transcript levels were rapidly upregulated. Upon injection of TAM into B16.F10 tumor-bearing *Trex1^FL/FL^R26^CreERT2^* mice, CD8^+^ T cells isolated from tumor tissue showed robust upregulation of the type I IFN-induced surface marker Sca-1, indicating IFN exposure (Fig. 1B). TREX1 iKO mice did not show any macroscopically detectable compromise of general health (not shown).

**Figure 1:**
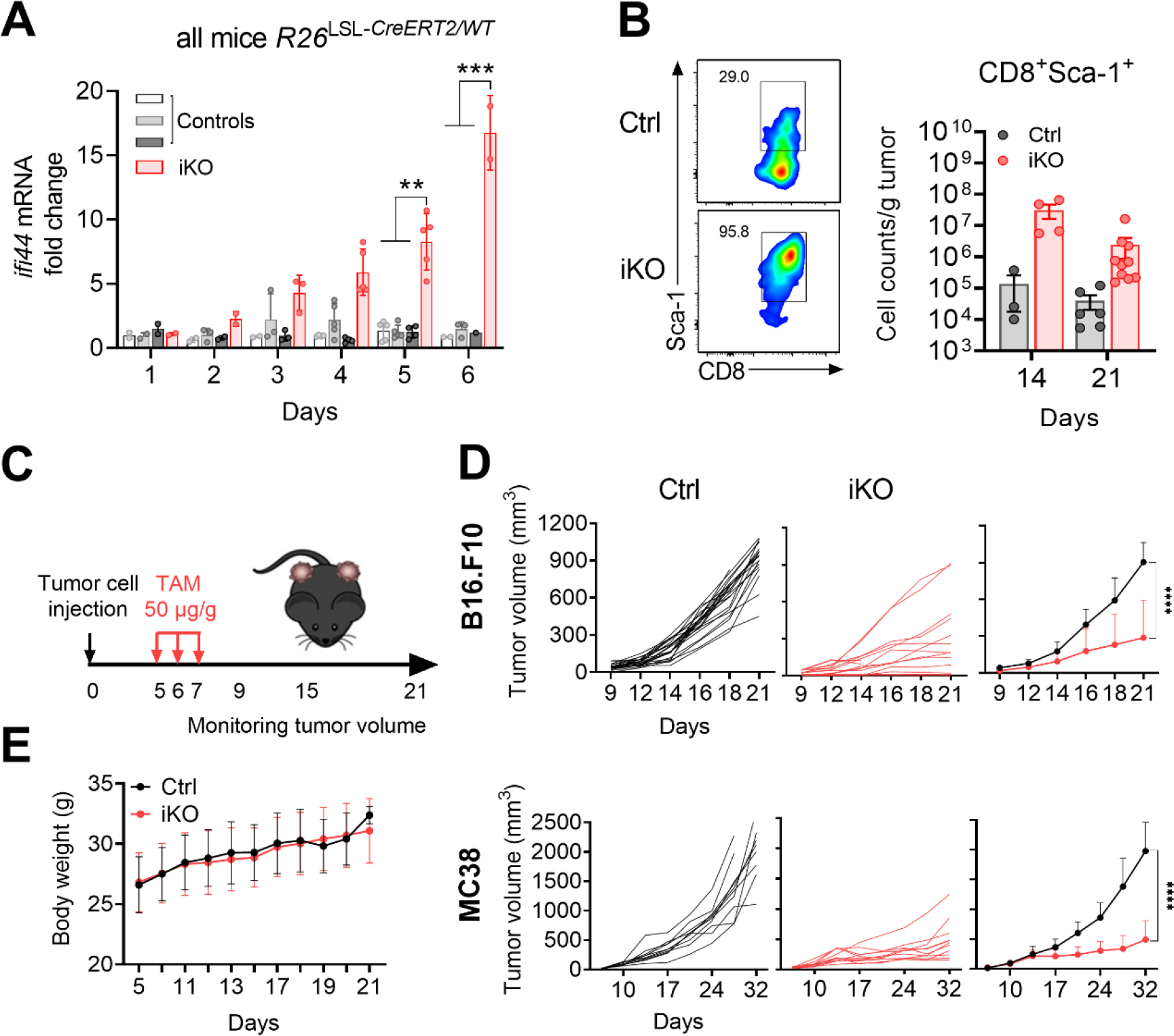
Systemic type I IFN response and improved tumor control in TREX1 iKO mice **A)** Ifi44 transcript levels (qRT-PCR) in total splenocytes at the indicated time points after induction of Trex1 inactivation in Trex1^FL/FL^R26^CreERT2^ mice by a single i.p. dose (1 mg) of TAM. Vehicle-treated Trex1^WT/FL^R26^CreERT2^ (white), TAM-treated Trex1^WT/FL^R26^CreERT2^ (light grey) and vehicle-treated Trex1^FL/FL^R26^CreERT2^ (dark grey) mice served as controls. Mixed effect analysis followed by Tukey’s multiple comparison test, means ± SD, **p<0.01*** p<0.001. **B)** Quantification of surface expression of SCA-1, encoded by the ISG Ly6a, on CD8^+^ T cells isolated from B16 tumors growing in Trex1^FL/FL^R26^CreERT2^ or Trex1^FL/WT^R26^CreERT2^ control hosts. All mice were treated with TAM at days 5, 6 and 7 and tumor tissue was harvested on day 21 after tumor inoculation. One-way ANOVA followed by Sidak’s multiple comparison test, means ± SD. **C)** Experimental schedule of induced Trex1 KO (‘TREX1 iKO’) in tumor bearing mice. **D)** Tumor growth in male TREX1 iKO or Trex1^FL/WT^R26^CreERT2^ control mice inoculated with 1×10^3^ B16.F10 cells intradermally on both hind limbs (upper panels) and in female mice inoculated with 3×10^5^ MC38 cells (lower panels). Each line in the left two graphs represents one tumor. Data are summarized in the graph on the right. N=18-24 tumors per group, means ± SD, two-way ANOVA, ****p<0.0001. **E)** Mean weight of tumor-bearing TREX1 iKO and control mice. N=15 per group, means ± SD.

To study the impact of TREX1 iKO in host cells on immune control of syngeneic tumors, we bilaterally implanted (*Trex1^WT^*) B16.F10 melanoma cells into *Trex1^FL/FL^R26^CreERT2^* and control mice by intradermal injection on the hind limb (Fig. 1C). Cre activity was induced by three injections of TAM on days 5, 6 and 7 after tumor cell implantation (Fig. 1C). In control mice, B16.F10 tumors grew rapidly, while tumor growth was much slower in TREX1 iKO mice, with day 21 tumor size reduced by about 70 % compared to control hosts (Fig. 1D). Again, neither hunching, alteration of feeding or social behaviour nor compromised weight gain were observed (Fig. 1E). Similar results were obtained with a second B16.F10 cohort (Fig. S1C), and using the MC38 colon adenocarcinoma (Fig. 1D) and MB49 bladder cancer models (Fig. S1D) Collectively, we show that induced inactivation of *Trex1* in non-malignant host cells triggers a systemic type I IFN response and significantly reduces growth of syngeneic tumors.

### Induced loss of TREX1 in host cells selectively inflames the tumor micromilieu but not other host tissues

To elucidate the impact of *Trex1* inactivation on local and systemic immune responses, we analysed immune cells from B16.F10 tumor tissue (Fig. 2A, S2A)) as well as tumor-draining LNs and spleens (Fig. S2B,C) from TREX1 iKO and control mice. Induced loss of TREX1 massively enhanced immune cell infiltration into the tumor. Numbers of CD45^+^ leukocytes were increased 10- to 100-fold compared to tumors of control hosts. This increase reflected enhanced infiltration by different cell types, including NK cells, DCs, and both CD4^+^ and CD8^+^ T cells. The increase in CD8^+^ T cells was also detected by immunofluorescence (Fig. 2B) and was more pronounced then the increase in CD4^+^ T cells, resulting in dominance of CD8^+^ over CD4^+^ T cells in tumors of TREX1-deficient hosts. Single-cell RNA sequencing of CD45^+^ cells FACS-sorted from cell suspensions of day 21 B16.F10 tumors (Fig. 2C, Fig. S3A) confirmed the increased CD8^+^-to-CD4^+^ T cell ratio in TREX1 iKO hosts (Fig. S3B) and revealed substantial alterations within the CD8^+^ T cell and the myeloid compartments. Cells responding to type I IFN in TREX1 iKO hosts were heterogeneously distributed, as determined by an ISG expression score which was primarily high in some subsets of monocyte/macrophages and a subset of T cells (Fig. S3C). Sizes of major immune cell populations in tumor-draining lymph nodes were largely unchanged except for an increase in DCs and NK cells (Fig. S2B). Slight increases in numbers of activated T cells were found in spleen (Fig. S2C). Flow cytometric analysis of tumors of the repeat B16.F10 cohort (Fig. S1C) yielded similar results (Fig. S2D) as did analysis of tumor-draining lymph nodes and spleen (not shown). In stark contrast to the robust immune cell infiltration of tumors, we did not observe inflammation in non-tumor tissues of the TREX1 iKO mice (Fig. 2D and S4) including heart and skeletal muscle, i.e. tissues that develop inflammatory changes in germline *Trex1^-/-^*mice.

**Figure 2:**
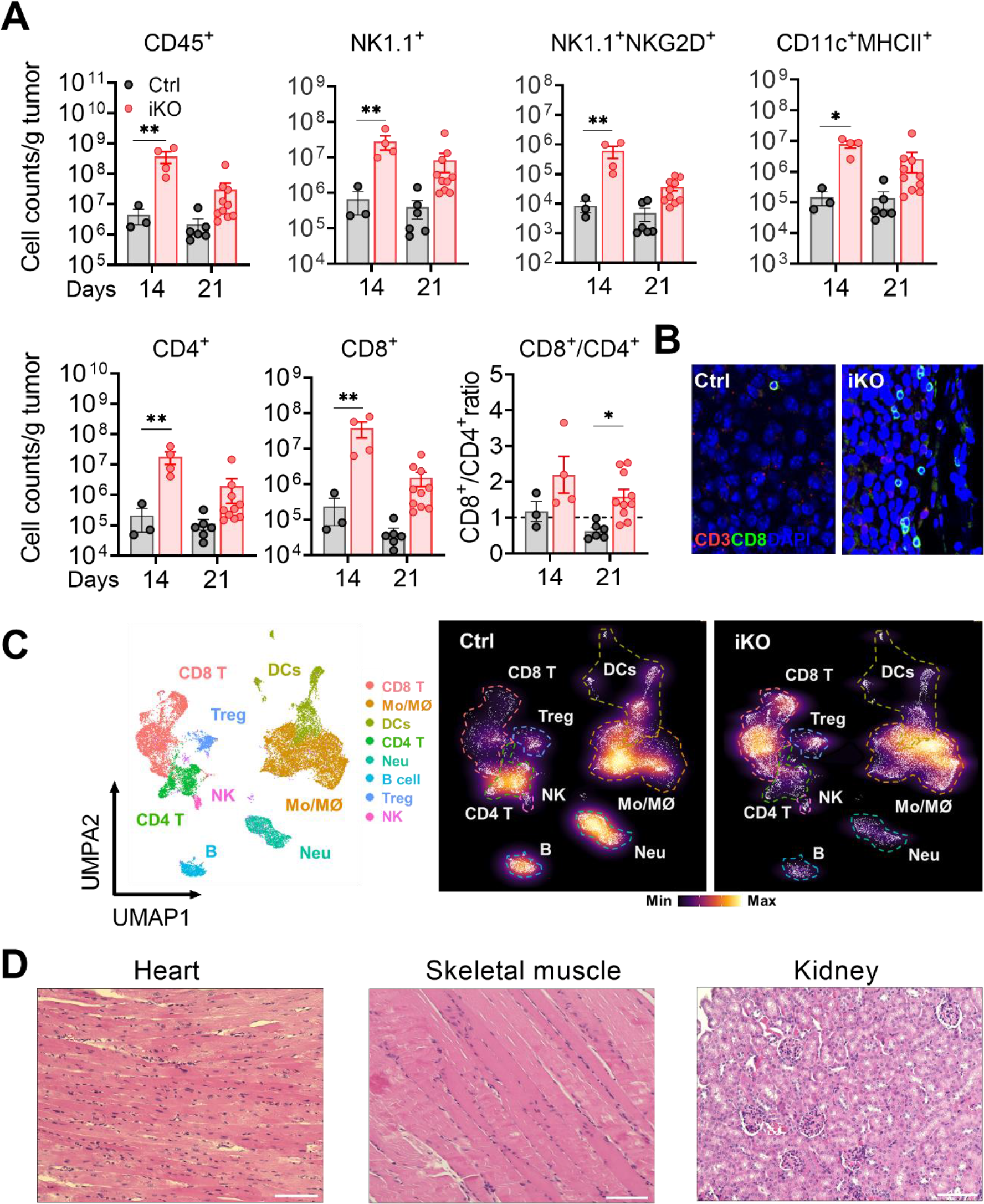
Induced loss of TREX1 in host cells potentiates immune infiltration of tumor tissue. A) Flow cytometric determination of absolute numbers of immune cells per gram of tissue in single cell suspensions of tumors harvested at the time points indicated. Immune cell populations were identified according to the gating shown in Fig. S2. Means ± SD, unpaired Student t test, *p<0.05, **p<0.01. **B)** Immunofluorescent staining of CD3 and CD8 on paraffin sections of tumors from control and TREX1 iKO hosts (representative of 3 mice/group). **C)** Single-cell RNA sequencing of CD45^+^ cells from tumors of TREX1 iKO or control hosts (N=4/group). Uniform manifold approximation and projection (UMAP) plot (left) of all 21,748 cells and density plots (right) showing relative frequencies of cells. **D)** H&E-stained paraffin sections of indicated tissues of TREX1 iKO mice sampled 5 weeks after TAM induction. Scale bars 100 µm.

Collectively, induced loss of TREX1 selectively inflamed the tumor microenvironment, with substantial increases of myeloid and T cell infiltration. Non-tumor tissues of the TREX1 iKO animals did not show signs of inflammation.

### Type I IFN- and CD8^+^ T cell-dependent tumor control upon induced loss of TREX1 in host cells

Immune stimulation occurring in the absence of TREX1 is entirely dependent on sensing of endogenous DNA via the cGAS/STING axis (26, 37–40). Activation of STING triggers production of type I IFNs by activation of IRF3, as well as NF-κB activation. Both, type I IFNs and NF-κB-dependent cytokines, such as TNF, mediate immune-stimulatory effects and could be responsible for enhanced anti-tumor immunity in TREX1 iKO mice, acting alone or synergistically. To test whether the type I IFN response of TREX1 iKO mice is essential for tumor control, we blocked the type I IFN receptor (IFNAR) by injection of anti-IFNAR1 antibody (Fig. 3A). Trex1 iKO mice featured a robust increase in surface expression of the ISG SCA1 by immune cells in tumor, tumor-draining LN and spleen, which was suppressed to control levels by IFNAR blockade (Fig. S5A). Loss of IFNAR signalling abrogated both, enhanced immune cell infiltration (Fig.3B) and improved immune control (Fig. 3C) of B16.F10 tumors in TREX1 iKO animals.

**Figure 3:**
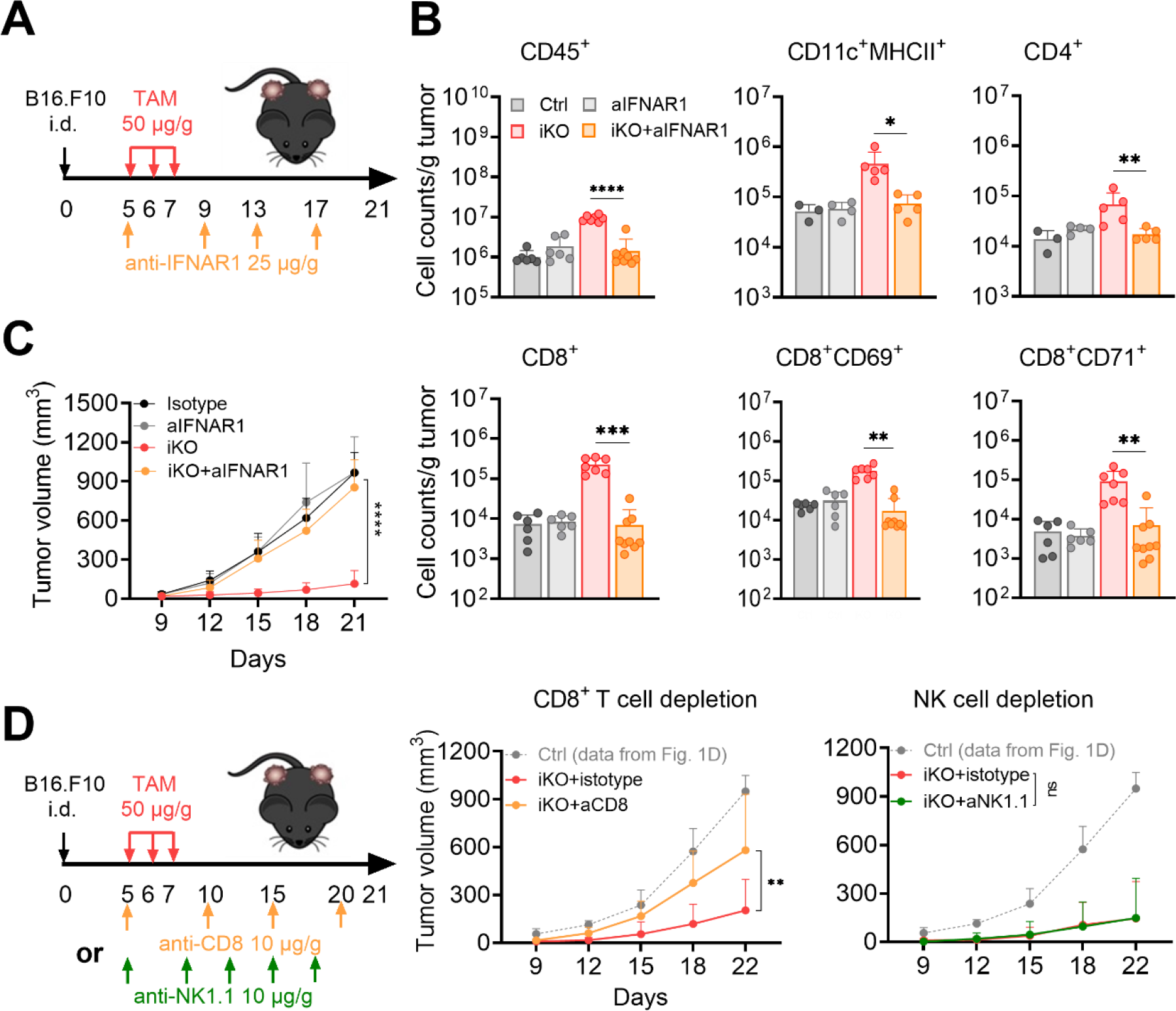
*Improved tumor control upon induced loss of TREX1 in host cells depends on type I IFN and CD8^+^ T cells*. ***A)*** Experimental schedule for analysis of the impact of IFNAR blockade on growth of B16.F10 tumors. ***B)*** Numbers of different immune cell types per gram of tumor tissue harvested on day 21. Means ± SD, One-way ANOVA followed by Sidak’s multiple comparison test, * p<0.05, ** p<0.01, *** p<0.001, **** p<0,0001. **C)** Tumor growth in TREX1 iKO and TAM-injected Trex1^FL/WT^R26^CreERT2^ control recipients that received injections of IFNAR-blocking antibody (anti-IFNAR1) or injections of isotype control antibody. N=5/per group, Means ± SD, two-way ANOVA (without post-hoc test), *** p<0.0001. **D)** Tumor growth in TREX1 iKO mice with or without injections of CD8^+^ T cell or NK cell-depleting antibody (tumor growth in TAM-injected Trex1^FL/WT^R26^CreERT2^ controls from Fig. 1D shown for comparison). N=5/per group, means ± SD, two-way ANOVA (without post-hoc test), *** p<0.001.

Numbers of both cytotoxic CD8^+^ T cells and NK cells were increased in tumors of TREX1 iKO hosts (Fig. 2A). To test whether one of these cell types was essential for tumor control, we used anti-CD8 or anti-NK1.1 antibodies to deplete them. In B16.F10 tumor-bearing TREX1 iKO mice, repeated injection of anti-CD8 (Fig. 3D) enabled efficient reduction of CD8^+^ T cells and accelerated tumor growth compared to isotype-treated TREX1 iKO mice, albeit not to the kinetics observed in TREX1-proficient hosts (Fig. 3D and S5B). While depletion of NK cells (Fig. 3D) was similarly efficient, loss of NK cells in tumor-bearing TREX1 iKO mice did not result in a detectable difference in tumor growth (Fig. 3D and S5C).

Collectively, type I IFN and CD8^+^ T cells, but not NK cells were essential for improved tumor control upon induced loss of TREX1.

### The intra-tumoral myeloid compartment undergoes inflammatory transformation with enhanced antigen presentation in TREX1 iKO hosts

For a higher resolution analysis of the TME in TREX1 iKO hosts, we selectively re-clustered the monocyte/macrophage and DC populations of the single-cell RNA seq experiment shown in Fig. 2C across control and TREX1 iKO tumor hosts (Fig. 4A and S6A). Within the monocyte/macrophage compartment, TREX1 iKO resulted in reduced fractions of cells expressing signatures of alternative activation or M2-like signatures (Fig. 4A cluster 1, Fig. S6C, J) or the signature of ‘angiogenic tumor infiltrating macrophages’ (angio-TAMs, as described by Ma et al. (41), Fig. S6D). This was accompanied by the appearance of a large macrophage fraction expressing M1-like signatures (Fig. 4A, cluster 0; Fig. 4B,J), commensurate with ‘IFN-stimulated-TAMs’ (41). These cells were largely absent in tumors of control hosts. Considering the much larger absolute size of the CD45^+^ population, M1-like cell numbers per volume of tumor tissue were increased by several orders of magnitude. These cells expressed the strongest ISG signature of all cell types of the TME (Fig. 4C) as well as the highest levels of CD40 (Fig. S6E). They also featured robust expression of genes involved in antigen presentation on MHC class I and MHC class II (Fig. 4E) including genes encoding components of the immunoproteasome (*Psmb10*, Fig. S6F; and others, not shown). Furthermore, these macrophages were marked by high transcript levels of the genes encoding the T cell attracting chemokines CXCL9 (Fig. S6G) and CXCL10 (Fig. 4D).

**Figure 4:**
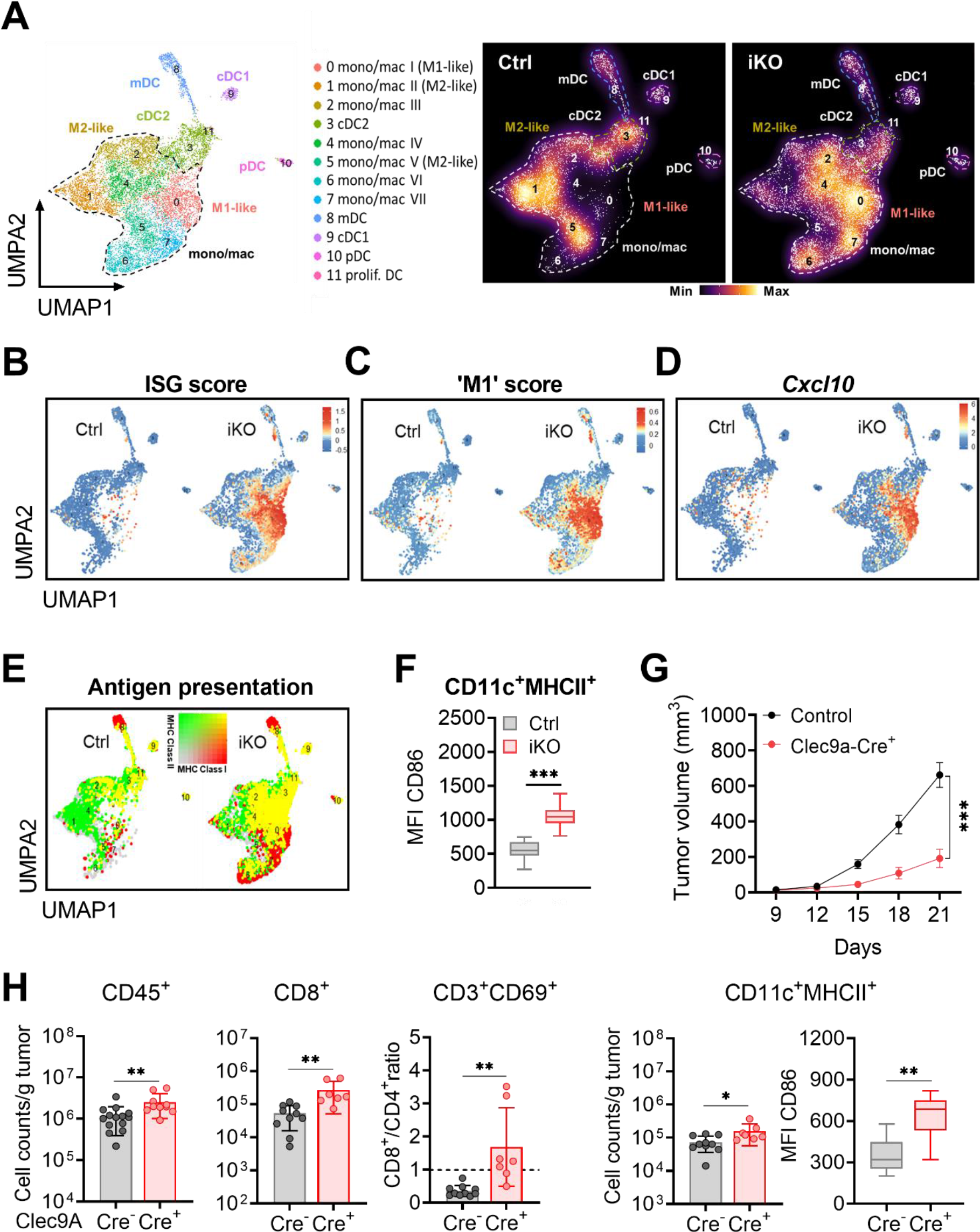
*Induced loss of TREX1 induces an inflammatory switch in the intra-tumoral myeloid compartment*. **A)** UMAP plot of myeloid cells across TREX1 iKO and control hosts (left) and density plots (right) representing frequencies of myeloid subpopulations in tumors of TREX1 iKO or control hosts. **B-D)** Feature plots showing expression of **B)** an IFN-induced transcriptional signature (ISG score was calculated from 40 ISGs, Table S1), **C)** a M1-like macrophage transcriptional signature, (score was calculated from 294 genes, Table S1), and **D** the Cxcl10 gene. **E)** Blended feature plot of expression of genes with functions in MHC class I or class II antigen presentation, (scores calculated from 13 genes each for MHC I and II-associated genes, Table S1). **F)** Mean expression of CD86 by tumor-infiltrating DCs in TREX1 iKO and control hosts determined by flow cytometry (from the experiment shown in Fig.2A) **G)** B16.F10 tumor growth in male Trex1^FL/FL^Clec9a-Cre^+^ or Trex1^FL/FL^ Cre-negative control mice. N=8/per group, means ± SD, two-way ANOVA without post-hoc test, **** p<0.0001. **H)** Flow cytometric determination of absolute numbers of immune cells per gram of tumor tissue in single cell suspensions of tumors harvested on day 21. Gating as in Fig. S2. Means ± SD, One-way ANOVA followed by Sidak’s multiple comparison test, * p<0.05, ** p<0.01.

DC populations did not show large changes in relative frequency in tumors of TREX1 iKO hosts, however, the absolute numbers of DCs per volume of tumor tissue increased 10-100 fold (see increase of total CD45^+^ cells and CD11c^+^MHCII^+^ cells as determined by flow cytometry, Fig. 2A and S2D). Single-cell RNA sequencing identified classical DCs (cDCs, *Zbtb46*^+^, Fig. 4A and S6A,B) clustered into cDC1 (*Clec9a^+^, Xcr1^+^, Itgae^+^,* cluster 9), cDC2 (*Itgam^+^, Sirpa^+^*, cluster 3) and mature/regulatory/migratory cDCs (mDC; *Ccr7^+^, Fscn1^+^, Cd200^+^, Cd80, Ly75^+^, Tnfrsf18^+^, Cd274^+^*, cluster 8), representing cells about to migrate to the tumor-draining lymph node (42) (Fig. 4A and S6A,B), as well as pDCs. Analysis of differentially expressed genes showed robust induction of ISG expression by all cDC populations in tumors of TREX1 iKO hosts (Fig. S6J, dwarfed in Fig. 4B by the even higher IFN response signature of monocyte/macrophage populations). CD8^+^ T cell-recruiting chemokines CXCL9 and CXCL10 were induced in TREX1 iKO cDC1 and mDC (Fig. S6J). Expression of genes essential for antigen presentation on MHC class I and class II was enhanced for all DC populations (Fig. 4E). Enhanced costimulatory capacity was reflected by increased mRNA levels of *Cd86* in mDCs (Fig. S6H) as well as increased surface CD86 expression by CD11c^+^MHCII^+^ cells as determined by flow cytometry (Fig. 4F and S2D).

Given these striking transcriptional signatures of enhanced antigen-presentation capacity of intratumoral DCs, we addressed whether TREX1 deficiency of only DCs improves tumor immune control. We had previously shown that constitutive absence of TREX1 selectively in cDCs of *Trex1^FL/FL^Clec9A-Cre* mice is sufficient to cause systemic autoimmunity (36, 43). Growth of transplanted tumors was robustly reduced in these animals and associated with increased intra-tumoral DC and CD8^+^ T cell numbers as well as enhanced DC expression of CD86 (Fig. 4G,H and S7). These findings suggest that cDC-intrinsic immune-stimulation significantly contributes to enhanced tumor control observed upon induced global *Trex1* inactivation.

Collectively, the massive increase in myeloid cells infiltrating tumors of TREX1iKO hosts was associated with striking changes toward more inflammatory phenotypes of monocytes and macrophages as well as enhanced antigen presentation capacity of monocytes, macrophages and DCs.

### Large increase of effector-like exhausted cells in the invigorated CD8^+^ T cell compartment of tumors in TREX1 iKO hosts

Enhanced CD8^+^ T cell-dependent tumor control in TREX1 iKO hosts was reflected in major alterations within the markedly expanded CD8^+^ T cell compartment. Selective unsupervised clustering of CD8^+^ T cells (across cells from control and iKO hosts, Fig. 5A and S8A, B) identified several clusters of cells expressing exhaustion signatures (variable mRNA levels of *Tox, Pdcd1, Lag3, Havcr2, Tigit, Entpd1,* clusters 0, 1, 4, 5, 6, 7, 10, 11, and 12). Unexhausted cells (clusters 2 and 3) comprised ‘naïve-like’/central memory cells (cluster 3: *Il7r^+^*, *Ccr7^+^, Sell^+^, S1pr1^+^, Tcf7^+^, Foxo1^+^, Lef1^+^*) as typically encountered in tumor tissue (44), as well as effector/effector memory cells (cluster 2: *Id2^+^, Ccl5^+^, Cxcr3^+^, Grzmk^+^, Fasl^+^*) variably expressing other effector-associated genes (*Zeb2, Tbx21, Klrg1, Grzma, Grzmb, Cx3cr1*) and memory-associated genes (*Slamf6, Eomes, Il7r, Tcf7, Lef1*). Some expression of *Tox* and *Pdcd1* in this cluster was not accompanied by significant expression of other co-inhibitory receptors. One cluster containing non-exhausted and exhausted cells expressed a robust signature of type I IFN-induced genes (including, *Isg20, Usp18, Rsad2, Ifit1, Ifit 3* and *Isg15*, among numerous other ISGs, cluster 8). Within the exhausted (T_EX_) compartment, a small population (lower part of cluster 7) identified as precursor exhausted cells (T_PEX_), was marked by signatures of ‘CD62L^+^ stem-like Tprex’, ‘CD62^-^ Tprex’ as described by Tsui et al. (45) or ‘T_EX_ progenitor 1’ as reported by Beltra et al. (46) (Fig. S8C, Table S1), including low, but detectable expression of *Ccr7, IL7r, Tcf7,* (Fig. S8C), as well as *Slamf6, Bcl6,* not shown. These cells expressed only intermediate levels of *Tox*, low levels of *Havcr2* and little T cell effector function (*Gzma, Gzmb, Gzmk, Prf1, Klrg1, Ifng* all low or negative). A large T_EX_ population featured a cell cycle signature (Fig. 5D) including e.g. *Mki67^+^ (Fig. S8B)* (clusters 4, 6, 9, 12). Several T_EX_ clusters variably co-expressed genes associated with effector function, including cytotoxicity (*Prf1, Gzmb*), *Ifng* and *Tnfrsf9* (Fig. 5D) (cluster 7 upper part, clusters 1, 5, 10, 11) equivalent to effector-like T_EX_ cells. Clusters 0 and 10 were characterized by highest levels of exhaustion-associated genes (*Tox*, coinhibitory receptors, *Entpd1*, *Prdm1*) and concomitant reduction of effector functions (lower expression of *Ifng* and cytotoxicity-associated genes) indicating differentiation toward terminal exhaustion (T_EX_term).

**Figure 5:**
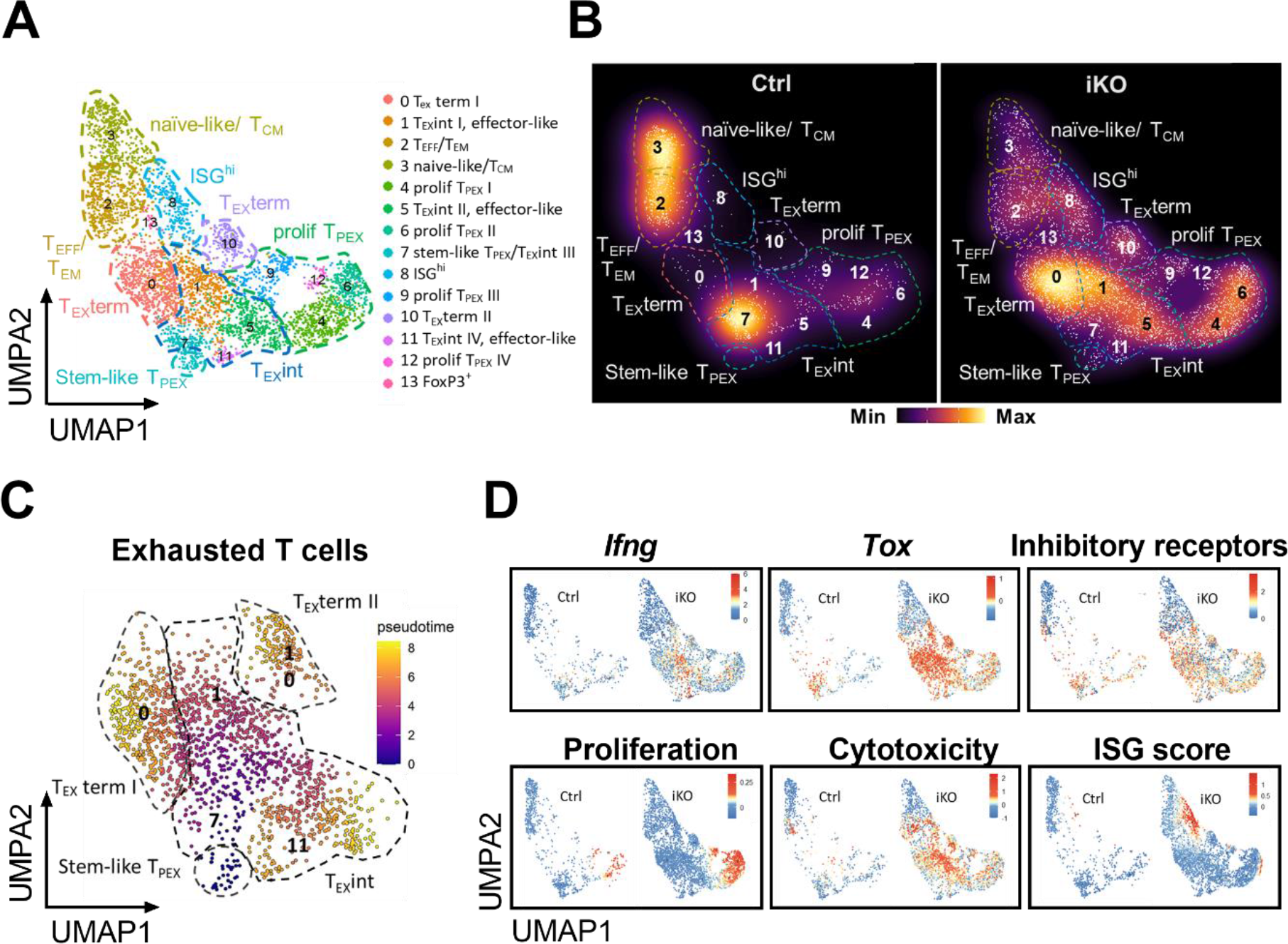
*Induced loss of TREX1 in host cells enhances infiltration of tumors with effector-like ‘exhausted’ CD8^+^ T cells*. **A)** UMAP plot of CD8^+^ T cells across TREX1 iKO and control hosts (left) and density plots **B)** representing frequencies of CD8^+^ T cell subpopulations in tumors of TREX1 iKO or control hosts. **C)** Supervised pseudotime analysis of (non-proliferating) T_EX_ populations using a single pre-defined root node (the stem-like T_EX_ progenitor subcluster of cluster 7). **D)** Feature plots showing expression of Ifng, Tox, and of co-inhibitory receptors (score calculated from 6 genes, Table S1), cell cycle-associated genes (score calculated from 185 genes, Table S1), cytotoxicity-associated genes (score calculated from 6 genes, Table S1), and an IFN-induced signature (score calculated from 40 ISGs, Table S1) in CD8^+^ T cells.

Loss of TREX1 in host cells profoundly changed the intra-tumoral CD8^+^ T cell response. Most importantly, the fractions of proliferating cells and cells with robust effector function showed a marked increase. The small CD8^+^ T cell population from tumors of control hosts (Fig. 2A) was dominated by naïve-like/memory populations, a small population of stem-like T_EX_ progenitors (cluster 7, lower part) and one T_EX_ population with moderate effector features (low or absent expression of *Ifng* and cytotoxicity-associated genes, Fig. 5A, B, cluster 7, upper part). In contrast, tumors of TREX1 iKO hosts contained vast numbers of cells co-expressing effector-associated genes (*Id2, Ifng, Prf1, Fasl, Grzmb,* and *Ccl3*, clusters 1, 5 and 11) along with the exhaustion signature. The small population of stem-like T_EX_ progenitors, 3.8% of total CD8^+^ T cells in control tumors (lower part of cluster 7), accounted for 1.0% of CD8^+^ T cells in tumors of TREX1 iKO hosts, reflecting about 2.5 to 25-fold higher absolute numbers, taking into account the 10- to 100-fold larger compartment size in tumors of TREX1 iKO hosts. Likewise, the fraction of proliferating intra-tumoral CD8^+^ T_EX_ cells (clusters 4, 6, 9 and 12) increased 1.7-fold in iKO compared to tumors of control hosts (Fig. S8D), corresponding to a 17- to 170-fold increase in absolute numbers per g of tumor tissue. In addition, fractions of cells differentiating toward terminal exhaustion (clusters 0 and 10), sparse in control hosts, were increased about 7-fold in iKO hosts (Fig. S8D), reflecting large increases of absolute numbers. Pseudotime analysis indicated differentiation trajectories from stem-like T_EX_ progenitors via effector-like T_EX_ cells toward terminal T_EX_ differentiation (Fig. 5C) in accordance with recent reports (reviewed in (47)). In addition to cells on the exhaustion trajectory, the absolute numbers of non-exhausted, cytolytic effector cells (part of cluster 2) also strongly increased in tumors of iKO hosts, despite showing reduced relative frequencies (Fig. 5A, D).

While tumors of control hosts contained few cells that strongly expressed a signature of type I IFN-induced genes, these cells formed a large population in TREX1 iKO tumors (cluster 8). A portion of these cells expressed memory-associated genes, whereas another expressed exhaustion features, prompting the question whether this cluster represents a distinct population defined by functional features beyond ISG expression. Subtraction of 40 strongly expressed ISGs and re-clustering of the CD8^+^ T cell compartment revealed that ISG^high^ cells now clustered with functionally very different populations, including naïve- and central memory-like, effector and effector memory cells, as well as most populations on the exhausted trajectory (Fig. S8E). Thus, expression of a prominent ISG signature, that was induced in a considerable fraction of T cells in TREX1 iKO hosts, is not a feature of one particular functional T cell subset, but rather occurs in functionally diverse CD8^+^ T cells.

In summary, we show that the increase in intra-tumoral CD8^+^ T cells is primarily accounted for by higher numbers of T_EX_ progenitors, effector-like and terminally exhausted T_EX_ cells.

### Synergy of induced TREX1 deficiency with checkpoint inhibition

The strongly increased numbers of CD8^+^ T cells differentiating along the exhaustion trajectory we found to be associated with improved tumor control in TREX1 iKO hosts suggested that immune protection may be further enhanced by concomitant immune checkpoint inhibition. As single cell transcriptome analysis had revealed increases in myeloid cells expressing *Cd274* (encoding PD-L1, Fig. 6A) and T cells expressing *Pdcd1* (encoding PD-1, Fig. 6B), we chose to combine induced inactivation of *Trex1* with PD-1 blocking antibody in B16.F10 tumor-bearing mice (Fig. 6C). While PD-1 blockade alone had little impact on tumor growth (Fig. 6D), the combination with TREX1 iKO resulted in efficient tumor control with complete rejection of 10 of 12 tumors of 6 mice, a result superior to all prior experiments with TREX1 iKO alone.

**Figure 6:**
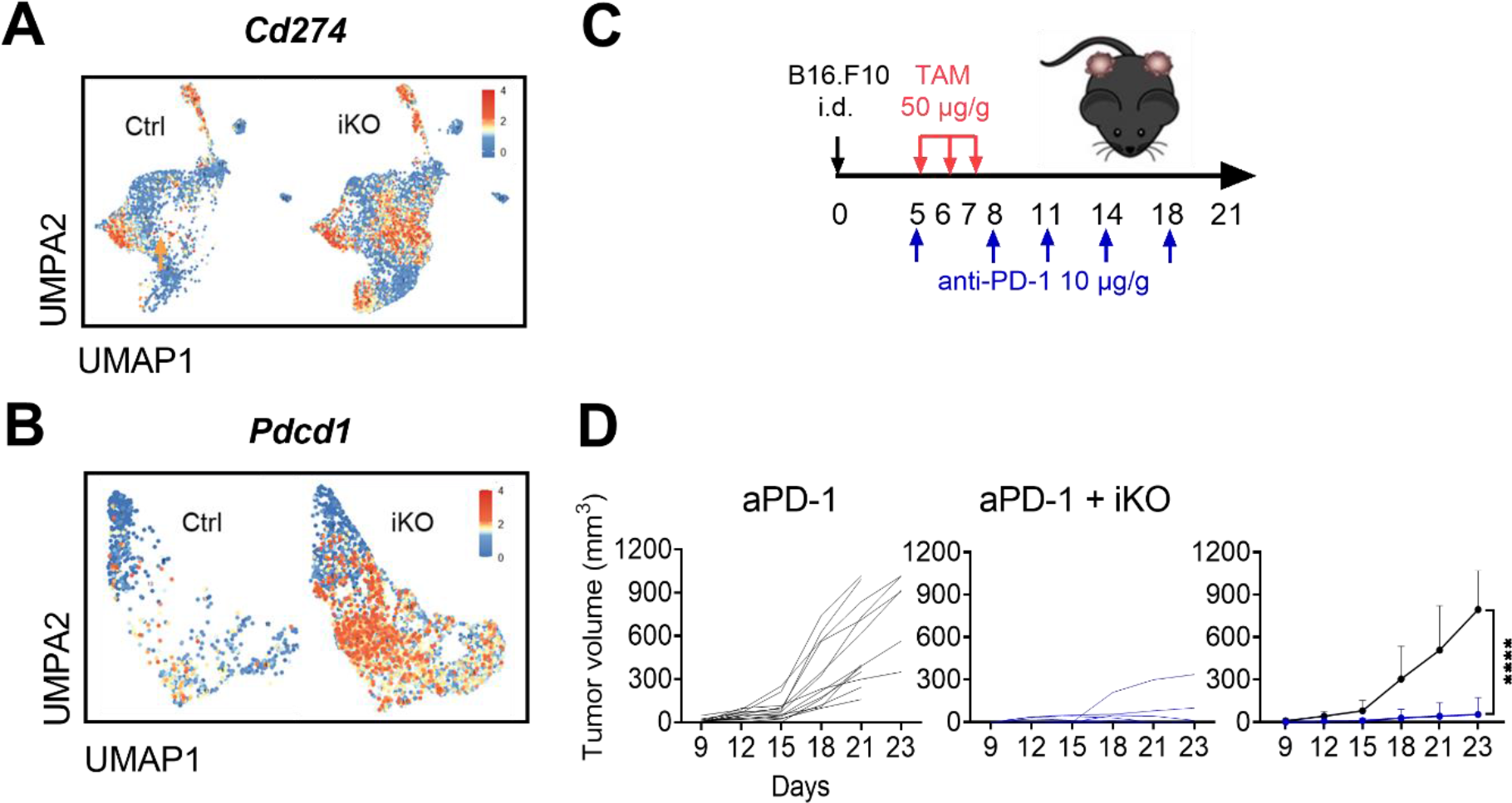
*Synergy of induced TREX1 deficiency with checkpoint inhibition*. **A-B)** Feature plot showing the intensity of expression of **(A)** Cd274, encoding PD-L1, in myeloid cells and **(B)** Pdcd1, encoding PD-1, in CD8^+^ T cells. **C)** Experimental schedule for induced TREX1 inactivation and anti-PD-1 administration in mice bearing B16.F10 tumors. Mice were inoculated with 1×10^3^ B16.F10 cells intradermally on both hind limbs. **D)** Tumor growth in male anti-PD-1-treated Trex1^FL/FL^R26^CreERT2^ and Trex1^FLWT^R26^CreERT2^ control mice. N=12 (6 mice) and N=14 (7 mice) tumors in TREX1 iKO and control hosts, respectively, Means ± SD, two-way ANOVA (without post-hoc test), **** p<0.0001.

Collectively, interference with host TREX1 activity synergizes with blockade of the PD-1 immune checkpoint.

### Cancer cell-intrinsic TREX1 deficiency synergizes with combined irradiation and checkpoint inhibition to induce long-term protection

The experiments in TREX iKO hosts tested the impact of induced TREX1 deficiency on the growth of TREX1-competent tumors. Three recent studies showed that lack of TREX1 in cancer cells enhances anti-tumor immunity and synergizes with PD-1 blockade (29–31). We tested control of established TREX1-deficient (*Trex1^-/-^*) B16.F10 tumors (Fig. S9) by treatment with combined immune checkpoint blockade and irradiation (Fig. 7A). Focal irradiation with 20 Gy slowed down the growth of control and *Trex1^-/-^* tumors (Fig. 7B). However, control of neither *Trex1* wildtype nor *Trex1^-/-^* tumors was improved compared to TREX1 competent tumors (Fig.7B, right). Combined irradiation and treatment with anti-PD-1 plus anti-CTLA4 was slightly more effective treating control and *Trex1^-/-^* tumors compared to irradiation alone (Fig. 7C). There was little difference in the response of control and *Trex1^-/-^* tumors to this treatment until about three weeks after tumor inoculation. However, longer observation revealed a profound effect of TREX1 deficiency. Whereas TREX1-proficient tumors, after a therapy-induced lag, progressed and reached large volumes 40 days post inoculation, all *Trex1^-/-^* tumors were completely rejected by four weeks after tumor inoculation. Re-challenge of these mice with WT B16.F10 cells resulted in no detectable tumor growth (Fig. 7C, far right), demonstrating induction of lasting memory. In summary, we confirm that TREX1-deficiency in cancer cells synergizes with combined irradiation and immune checkpoint inhibition to enhance stimulation of anti-tumor immunity and demonstrate formation of lasting protective memory.

**Figure 7:**
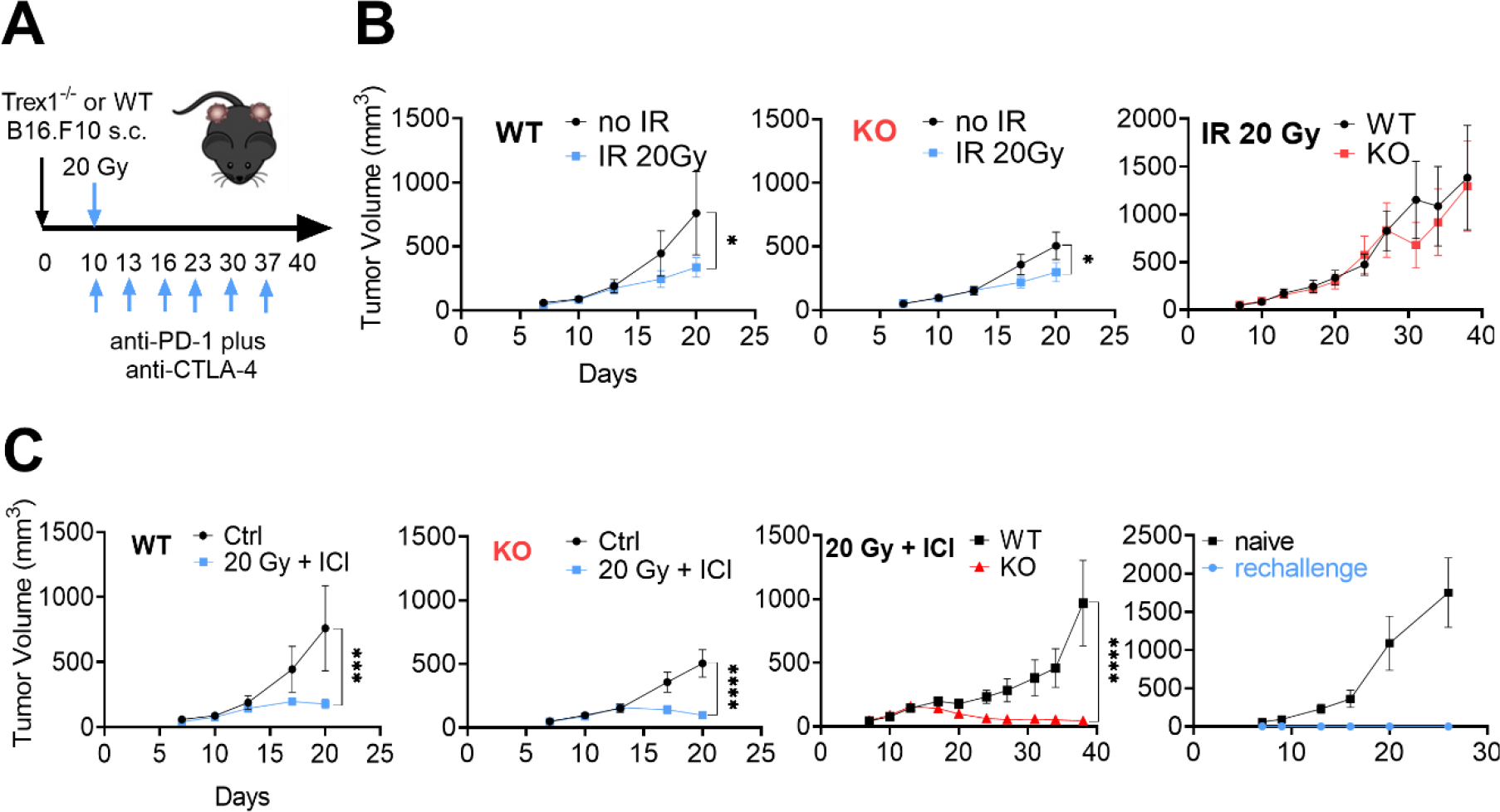
*Constitutive TREX1 deficiency in cancer cells combined with irradiation and checkpoint inhibitors enables long-term protective immunity*. **A)** Experimental schedule for analysis of Trex1^-/-^ (KO) or wildtype (WT) B16.F10 tumor growth in wild-type males. **B)** Half of the mice bearing wildt-type or KO tumors were locally irradiated with 20 Gy. N=8, means ± SD, two-way ANOVA (without post-hoc test), **** p<0.0001. Irradiated WT and KO tumors were monitored for 40 days. **C)** Half of the mice bearing WT or KO tumors received treatment with local irradiation (20 Gy) plus combined anti-CTLA-4 and anti-PD-1 checkpoint inhibitors (ICI). N=8, means ± SD, two-way ANOVA (without post-hoc test), **** p<0.0001. Irradiated and checkpoint inhibitor-treated WT and KO tumors were monitored for 40 days. Tumor growth after re-challenge of the irradiated plus ICI-treated KO mice with WT B16.F10 cells 4 weeks after complete regression of their tumor. N=4 per group, 1 tumor per mouse, means ± SD, two-way ANOVA (without post-hoc test), **** p<0.0001.

## Discussion

Overcoming the immune-suppressive tumor microenvironment of ‘cold’ tumors, thereby sensitizing them for therapy with immune checkpoint blockers, has emerged as a key task of immuno-oncology. Pharmacological inhibitors of TREX1 for interference with cytosolic DNA disposal and activation of cell-intrinsic cGAS/STING responses in cancer tissue are currently an intense focus of research. While much effort was devoted to evoking immune-stimulation by interference with cancer cell-expressed TREX1 (29–31), we show that, even if restricted to non-malignant host cells, loss of cytosolic DNA disposal by induced inactivation of TREX1 results in robust inflammation of the tumor microenvironment, invigorates the CD8^+^ T cell response and notably improves immune control of even largely immunotherapy-resistant syngeneic tumors. Tumor immune control upon TREX1 iKO was dependent on type I IFN and CD8^+^ T cells and synergized with ICI. Partly due to their genetic instability, cancer cells accumulate large amounts of different species of endogenous DNA in their cytoplasm that can engage cell-intrinsic cGAS. Our data show that, upon induced loss of TREX1 in adult animals, steady-state levels of endogenous immune-stimulatory cytosolic DNA in non-malignant cells, including immune cells, is sufficient to drive efficient CD8^+^ T cell-mediated tumor immune control. This is advantageous as intra-tumoral cGAS/STING activation may not always be an option and is often compromised by mutation, epigenetic silencing or other mechanisms as a means of cancer immune evasion (32–35), a problem avoided by systemic TREX1 inactivation in non-malignant host cells.

Importantly, acute inflammation upon induced loss of TREX1 selectively occurred in the tumor but spared other (non-tumor) tissues of the TREX1 iKO animals. In particular, heart and skeletal muscle, organs that are heavily inflamed in constitutive *Trex1^-/-^* mice (26, 48) did not show any alteration on histology and the animals did not develop any macroscopic signs of disease. In notable contrast, induced deletion of another AGS-causing gene in adult animals, *Adar1*, results in lethal inflammation within days (49). Induced systemic TREX1 deficiency may achieve selective acute inflammation of tumors, but not other organs, due to local priming of inflammatory signaling pathways in the tumor tissue that are mostly, but not completely suppressed by the anti-inflammatory micro-milieu. This finding suggests that therapeutic windows of treatment duration and dose may exist for systemic pharmacologic interference with TREX1 activity in cancer patients, enabling generation of a sufficient adjuvant effect to boost anti-tumor immunity while avoiding induction of severe autoimmunity. Of note, immune stimulation by induced Trex1 inactivation differs from the phenotype of germline-encoded TREX1 deficiency. Development of the immune system under conditions of chronic cGAS/STING stimulation often causes inflammatory pathology at *in utero* stages of ontogeny in patients and mice, albeit with considerable variation, whereas induced systemic loss of TREX1 seems to be well tolerated for significant amounts of time by adult mice.

Inflammation of the TME triggered by induced loss of TREX1 in host cells entailed robust increases of absolute numbers per tumor volume of monocytes and macrophages, conventional and plasmacytoid DCs, NK cells as well as CD4^+^ and CD8^+^ T cells compared to tumors of control hosts. Regulatory M2- like macrophage populations, key determinants of an immune-suppressive, tumor-promoting micro- milieu dominated control tumors and were largely replaced by inflammatory M1-like cells expressing high levels of CD40 and signatures of antigen presentation in TREX1-deficient hosts.

The well-documented essential functions of DCs in T cell-mediated anti-tumor immunity are largely dependent on type I IFN responses mounted by DCs themselves (50, 51). cDC1 play key roles in cross-presentation of tumor antigen, priming of tumor-specific CD8^+^ T cells and maintaining stem-like CD8^+^ T_PEX_ cells in tumor-draining lymph nodes (52–54). cDC1 also recruit CD8^+^ T cells into the tumor and re-stimulate them locally, thereby promoting their proliferation, effector function and survival (52, 53). To enable crucial intra-tumoral cDC1/CD8^+^ T cell interactions, cDC1 attract T cells by secretion of CXCL9 and CXCL10 (55). STING agonism can enhance DC function (5); however, in some contexts it can eliminate cDC1s (56). Systemic STING activation resulting from induced loss of TREX1, increased absolute intra-tumoral numbers of *Ccr7*-negative cDC1 and cDC2, as well as *Ccr7*-expressing mature/migratory DCs. All three subsets upregulated transcriptional signatures of the type I IFN response as well as antigen presentation and co-stimulation. Thus, systemic inactivation of *Trex1* in host cells represents a mode of STING activation that not only avoids loss of DC populations, but on the contrary increased their numbers 10- to 100-fold. MHCII^+^ CD11c^+^ DCs were also moderately increased in the tumor-draining LN and remained unchanged in spleen. Our observation of, on the one hand, higher numbers of *Ccr7*-negative cDC1s, expressing *Cxcl9* and *Cxcl10*, and exhibiting upregulated signatures of antigen presentation, and on the other hand, higher numbers of stem-like T_PEX_ and much increased proliferating CD8^+^ T cells compared to tumors of control host, are compatible with enhanced intra-tumoral CD8^+^ T cell restimulation by cDC1 upon induced loss of TREX1. An important role of cDCs in the improved tumor control of TREX1 iKO mice was corroborated by similar protection observed in mice with cDC-specific (albeit constitutive) *Trex1* inactivation.

T cell-mediated protective immunity against chronically persisting antigen, as in malignant disease, depends on supply of effector-like ‘exhausted’ (T_EX_) cells originating from largely lymph node- dwelling stem-like progenitors of exhausted CD8^+^ T (T_PEX_) cells (47). The exhaustion program enables these T_EX_ cells to survive and maintain significant levels of effector function despite chronic TCR over- stimulation (47). PD-1 blockade enhances the anti-tumor T cell response by promoting T_PEX_ proliferation and differentiation of their progeny along the exhaustion trajectory (46, 47, 57). We herein found that induced loss of TREX1 represents an alternative principle to invigorate effector-like T_EX_ generation. The dramatic increase of absolute numbers of CD8^+^ T cells in tumors of TREX1 iKO hosts was primarily accounted for by expanded populations of the T cell exhaustion trajectory (46). T_PEX_ cells expressing *Tcf7* and *Ccr7,* which are primarily confined to tumor-draining LNs (45, 47, 58), were also represented as a small fraction in the B16 tumor tissue, as reported previously for melanoma and some other tumors (55, 58). Induced loss of TREX1 increased their absolute numbers and also massively increased numbers of proliferating, Tcf7-negative T_PEX_ (46). This large proliferating T_PEX_ pool gave rise to large numbers of effector-like T_EX_ cells. These cells, largely absent in control tumors, expressed high levels of IFN and cytotoxic molecules, including granzymes and perforin, and likely mediated tumor control in the TREX1 iKO hosts. At the endpoint of the T_EX_ differentiation path were vast numbers of cells expressing signatures of terminal exhaustion with high levels of co-inhihitory receptor mRNA in tumors of TREX1 iKO hosts. The massive proliferative activity of T_PEX_ that we observed, together with the firmly established short lifespan of terminally exhausted CD8^+^ T cells (46) indicates highly dynamic turnover of cells along the T_EX_ trajectory, driven by the continuous STING response in the TREX1- deficient immune system, replenishing cells with robust effector function. TREX1 iKO also expanded a small population of non-exhausted T_EFF_ cells. Our data do not allow for an assessment of relative contributions of these T_EFF_ cells versus the much larger compartment of effector-like T_EX_ cells to tumor immune control. We also did not address the relative contribution of myeloid cell- versus T cell-intrinsic STING activation to the improved tumor control upon TREX1 iKO. T cell-intrinsic cGAS engagement by endogenous DNA and autocrine type I IFN effects were reported essential for efficient T cell-mediated tumor control (59). High STING signaling intensity in CD8^+^ T cells may result in compromised proliferation and cell death (16, 17, 60). In contrast, loss of TREX1 results in only mild STING activation, which is known to be blunted by haploinsufficiency of cGAS (39, 40) or STING (26). This might enable enhanced CD8^+^ T cell-mediated immunity, with important implications for immuno-oncology and cellular therapy approaches. Furthermore, we previously showed that deletion of *Trex1* selectively in B cells does not lead to STING activation or detectable transcriptional alterations, while they retain responsiveness to STING agonists (36). Hence the problem of differentiation of large numbers of immunosuppressive IL-35^+^ Breg cells limiting immune activation in studies using STING agonist treatment (23) should not occur in STING activation through systemic induced loss of TREX. Indeed, the B cell fraction decreased in tumors of TREX1 iKO hosts and little transcriptional changes were observed except for very a mild ISG signature with no evidence of Breg cell differentiation (not shown).

As tumor immune control in Trex1 iKO animals was associated with robust induction of PD-L1 on myeloid cells along with large increases in CD8^+^ T cells expressing PD-1 and other IRs, we expected that TREX1 iKO should synergize with immune checkpoint inhibition. Indeed, additional PD-1 blockade initiated simultaneously with tamoxifen injections for TREX1 iKO induction resulted in further improvement of tumor immune control with complete rejection of most tumors. This finding suggests that inactivation of TREX1 sensitizes the tumor for immune checkpoint inhibitor therapy. In human cancer, pre-treatment with a TREX1 inhibitor followed by checkpoint inhibition only few weeks later (after STING-induced inflammation of the TME occurs) may constitute an efficient treatment regime.

Cancer cell-intrinsic TREX1 is an important counter-regulator of anti-tumor immunity. Therapy-induced genomic damage enhances cytosolic DNA accumulation (9) and cGAS/STING-induced upregulation of TREX1 (28, 31). Interfering with cancer cell-expressed TREX1 can potently stimulate tumor immune control (29–31). Accordingly, we observed that upon combined irradiation and ICI, deficiency for TREX1 in tumor cells enabled complete tumor rejection and generation of protective long-term memory. As efficient, selective inhibition of cancer cell TREX1 activity may not always be possible and since inactivation of the cGAS/STING axis enables immune evasion in many cancers, our finding that induced loss of TREX1 in non-malignant host cells alone efficiently stimulates anti-tumor immunity greatly improves the prospect of a successful clinical translation of TREX1 inhibition for cancer immunotherapy.

Important open questions include the impact of induced systemic TREX1 inactivation on lymph node- dwelling T_PEX_ cells and on the durability of the invigorated T cell response and memory induction compared to ICI alone. Moreover, we are currently investigating the impact of TREX1 iKO on formation of intra-tumoral cDC1/CD8^+^ T cell clusters ref (55).

Collectively, we show that systemic targeting of *Trex1* in host cells, resulting in global low-level STING activation, induced inflammation selectively in the tumor microenvironment and enhanced T cell-mediated tumor control that synergized with checkpoint inhibition, suggesting important therapeutic potential for pharmacological systemic interference with TREX1 nuclease activity. Inflammation of the TME, but not non-tumor tissues, indicates that therapeutic windows of dose and treatment duration may enable efficient ‘heating up’ of human tumors while avoiding autoimmunity. Our data may inform the development of TREX1 inhibitors with respect to pharmacokinetics. TREX1 nuclease inhibitors have been reported in different stages of development (61). The principle of systemic therapeutic inhibition of TREX1 activity may be of relevance not only for enhancing endogenous anti-cancer T cell responses, but also for treatment with engineered tumor-antigen-specific T cells and as an adjuvant principle for vaccination. TREX1 inhibition might also boost immunity in chronic infection.

## Material & Methods

### Mice

*Trex1^FL/FL^Rosa26^CreERT2^* and *Trex1^FL/FL^Clec9a-Cre* mice were described earlier (36). Mice were housed under specific pathogen-free conditions at the Experimental Center, Medical Faculty Carl Gustav Carus, TU Dresden, the Interfaculty Biomedical Facility, University of Heidelberg, and the Bristol Myers Squibb, Redwood City, CA. All procedures were in accordance with institutional guidelines on animal welfare and were approved by the Landesdirektion Dresden, Regierungspräsidium Karlsruhe or the Institutional Animal Care and Use Committee at Bristol Myers Squibb.

### Cell lines

Mouse melanoma and adenocarcinoma cell lines B16-F10 and MC38 were purchased from ATCC. Mouse bladder carcinoma cell line MB49 was kindly provided by the Urology Department University Hospital, Medical Faculty Carl Gustav Carus Dresden. B16-F10 cells were cultured in DMEM media (Gibco, Waltham, MA) supplemented with 10% heat inactivated FCS (Biochrom GmbH, Berlin, Germany), nonessential amino acids (Biochrom GmbH), 100 U/ml penicillin (Biochrom GmbH) and 100 μg/ml streptomycin (Biochrom GmbH). MB49 was cultured in RPMI media (Biochrom GmbH) supplemented with 10% FCS (Biochrom GmbH) and 50 μg/ml Gentamycin (Sigma, Darmstadt, Germany).

### Generation of TREX1-deficient B16.F10 cells

Ribonucleoprotein-based CRISPR mutagenesis was performed on parental B16.F10 cells using one of 3 different guide RNAs, including guide 3 (5’-GGC.UCA.CAG.ACC.CUG.CCC.CA-3’). Polyclonal gene-edited populations were subcloned and clones were analyzed for *Trex1* inactivation by sequencing and western blot. Trex1 KO clone #3-2, originating from targeting using guide 3 was used for *in vivo* experiments.

### *In vivo* analysis of tumor growth

Eight-to-12 week-old male and female *Trex1^Fl/Fl^ R26^CreERT2^* mice were used. A total of 1×10^3^ *Trex1* wildtype B16-F10 cells or 2.5×10^5^ MB49 cells were mixed with an equal volume of Matrigel (BD Biosciences, Franklin Lakes, NJ) and intradermally injected on both hind limbs of male mice on day 0. A total of 3×10^5^ MC38 cells were injected intradermally. Mice received 1 mg of tamoxifen (Sigma Aldrich, St. Louis, MO) in 100 µl sunflower oil/5% ethanol by i.p. injection on day 5 for three consecutive days.

For *in vivo* immune checkpoint blockade experiments, 200 µg anti-PD-1 (clone RMP1-14, BioXcell, Lebanon, NH) or rat IgG2b isotype (clone 2A3, BioXCell) in 200 µL PBS was injected i.p. on days 9, 12, 15, 18 and 21. For depletion experiments, anti-CD8 (clone 2.43, BioXcell, 200 μg in 200 µl of PBS) or anti-NK1.1 (clone PK136, Bioxcell, 300 μg in 200 ul of PBS) antibodies and corresponding isotype controls rat IgG2b (clone LTF2, BioXcell) and mouse IgG2b (clone C1.18.4, BioXcell), respectively, were administrated i.p. on days 5, 10, 15 and 20. For *in vivo* IFNAR1 blockade, 500 μg in 200 µl PBS of anti-IFNAR1 antibody (MAR1-5A3, BioXcell) and isotype control (MOPC-21, BioXcell) were injected i.p. on days 5, 9, 13, 17 and 21. Perpendicular tumor diameters were measured using a caliper. Tumor volume was calculated as L × W^2^/2, where L is the longest diameter and W the perpendicular diameter.

For analysis of *Trex1^-/-^* B16.F10 tumor growth, 5×10^5^ B16.F10 clone 3-2 or control cells were injected subcutaneously into wild-type mice on day 0. On day 10, the tumor size was approximately 80-90 mm^3^ and checkpoint inhibition was initiated by injection of anti-PD1 (clone 4H2) plus anti-CTLA4 (clone 9D9); both antibodies were administered at 10 mg/kg body weight. The same doses of KLH-IgG1 (D265A) and IgG2a were injected as isotype controls. Mice that received irradiation were given a single 20 Gy dose using a Small Animal Radiation Research Platform (SARRP, Xstrahl, Inc.) on the first day of ICI dosing.

### Preparation of single-cell suspensions for flow cytometry

Tumors, tumor-draining lymph nodes and spleens were harvested at day 14 or 21 post-tumor implantation. Tumors were minced and digested with an enzyme mixture containing Liberase (25 µg/ml, Liberase^TM^ Research Grade, Sigma Aldrich), Hyaluronidase (0.5 mg/ml from bovine testes, Sigma Aldrich), and DNaseI (200U/µl, grade II, from bovine pancreas, Sigma Aldrich) in DMEM/20mM Hepes buffer for 30 min at 37°C with agitation (1300rpm). Digested tumor tissue was minced, passed through a 100 µm filter, then washed with 5 ml cold FACS buffer (0.5% BSA/2mM EDTA/PBS), and centrifuged at 1200 rpm for 5 min at 4°C. CD45^+^ cells were enriched from bulk tumor cell suspensions using CD45 (TIL) mouse Microbeads kit followed by magnetic separation on LS columns (Miltenyi Biotec, Bergisch-Gladbach, Germany). After eluting CD45^+^ cells from the column, cells were pelleted by centrifugation at 1200 rpm for 5 min at 4°C. Lymph nodes and spleens were minced, passed through a 40 µm mesh and centrifuged at 1200 rpm for 5 min at 4°C. Cell pellets were resuspended in 100 µL of Fc-blocking medium (5 µg/ml Purified anti-mouse CD16/32 Antibody (Biolegend, San Diego, CA) in FACS buffer). After a 20 min incubation on ice, 100 µl FACS buffer containing antibodies was added. The staining mixtures were incubated for another 45 min on ice. After washing, pelleted stained cells were resuspended in FACS buffer containing 1 µg/ml DAPI and kept on ice until analysis.

### Staining for flow cytometry

Antibodies used included anti-mouse CD45 (PerCP-Cy5.5 conjugated, clone 30-F11, eBioscience, Waltham, MA), anti-mouse CD3 (FITC conjugated, clone 17A2, Biolegend), anti-mouse CD4 (PE/Cy7 conjugated, GK1.5, eBioscience), anti-mouse CD8 (APC/Cy7 conjugated, clone 53-6.7 Biolegend), anti-mouse NK1.1 (APC conjugated, clone PK136, Biolegend), anti-mouse NKG2D (PE conjugated, clone 145-2C11, eBioscience), anti-mouse Ly-6A/E (Sca-1) (V500 conjugated, clone D7, BD Bioscience) anti-mouse CD11b (PE conjugated, clone M1/70, eBioscience), anti-mouse CD11c (APC conjugated, clone N418, Biolegend), anti-mouse MHCII (FITC conjugated, clone M5/114.5.2, eBioscience), anti-mouse CD86 (APC/Cy7 conjugated, clone GL1, Biolegend), anti-mouse CD3 (biotin conjugated, clone 145-2C11, eBioscience), anti-mouse CD19 (biotin conjugated, clone eBio1D3, eBioscience), anti-mouse NK1.1 (biotin conjugated, clone PK136 eBioscience), anti-mouse F4/80 (biotin conjugated, clone BM8, eBioscience), anti-mouse CD3 (APC conjugate, clone 145-2C11, eBioscience), anti-mouse CD44 (APC/Cy7 conjugated, clone GL1, Biolegend), anti-mouse CD62L (BV510 conjugated, clone MEL14, Biolegend), anti-mouse CD69 (PE conjugated, clone H1.2F3, Biolegend), anti-mouse CD71 (eFluor 450 conjugated, clone R17217, eBioscience). SA-V500 (BD Biosciences) was used for detection of biotin-conjugated antibodies. All data acquisition was done using a FACSAria III (BD Bioscience) and analyzed using FlowJo software (FlowJo LLC, Ashland, OR).

### Single-cell RNA sequencing

Preparation of single cell suspensions from tumor tissue and staining for extracellular markers were performed as described above for flow cytometry. Upon digestion of tumor tissue from 4 control and 4 TREX1 iKO animals followed by CD45^+^ cell enrichment, cells were resuspended in 50 µL Fc blocking medium (2.5 µg/ml purified anti-mouse CD16/32 Antibody (BioLegend) in Cell Staining Buffer (BioLegend)). Individual samples were barcoded by incubation with TotalSeq^TM^ antibodies (BioLegend, San Diego, CA) for 30 min on ice. After multiplexing, cells were pooled and stained with anti-mouse CD45 (PerCP-Cy5.5-conjugated, clone 30-F11, eBioscience) antibody. Samples were sorted on a FACSAria III instrument (BD Bioscience) by gating on live, CD45^+^ single cells.

Droplet-based scRNA-seq was performed using the 10x Genomics Chromium Single Cell Kit v3.1 An aliquot of the single cell suspension was visually inspected under a light microscope to check viability and cell density. Single-cell suspensions of 1-2×10^3^ cells per microliter with viability of more than 90% were mixed with reverse transcription mix and nuclease-free water according to the Chromium manual, aiming for 30 ×10^3^ cells per reaction. Cells were loaded on a Chromium Single Cell G Chip on the 10x Genomics Chromium system. In short, the droplets were directly subjected to reverse transcription, the emulsion was broken and cDNA was purified using silane beads. After amplification of cDNA (10 cycles) using primers to enrich cDNA as well as Totalseq-A hashtags, the samples underwent SPRI bead purification and fractionation for smaller fragments (up to 400 bp) to enrich the hashtag sequences and larger fragments (>400 bp) to enrich cDNA fragments. After quality check and quantification, the 10x Genomics single cell RNA-seq library preparation, involving fragmentation, dA-tailing, adapter ligation and 12-cycle indexing PCR, was performed based on the manufacturer’s protocol. In parallel, the hashtag library was prepared by a 10-cycles indexing PCR. After quantification, both libraries were sequenced on Illumina NovaSeq 6000 S4 flow cells in 200 bp paired-end mode, generating 1600-1800×10^6^ fragment pairs for the gene expression libraries and a minimum of 70×10^6^ fragment pairs for the hashtag library. Sequencing data are available under the GEO accession number GSE268792.

### Single-cell data analysis

The raw sequencing data was then processed with the ‘count’ command of the Cell Ranger software (v- 7.0.0) provided by 10X Genomics with the option ‘--expect-cells’ set to 10,000 (all other options were used as per default). To build the reference for Cell Ranger, mouse genome (GRCm39) as well as gene annotation (Ensembl 104) were downloaded from Ensembl and the annotation was filtered with the ‘mkgtf’ command of Cell Ranger (options: ‘--attribute=gene_biotype:protein_coding -- attribute=gene_biotype:lincRNA –attribute=gene_biotype:antisense’). Genome sequence and filtered annotation were then used as input to the ‘mkref’ command of Cell Ranger to build the appropriate Cellranger Reference. Matrix was processed for downstream data analysis using the R program based on the Seurat Package. The quality control threshold was set as less than 5% of mitochondrial RNA and nFeature RNA greater than 200. Gene score, marker genes of each cluster, and ISG regression were generated using the Seurat package according to manufactory. Trajectory analysis was performed with Monocle3.

### Histology

Sections (5 µm) of formalin-fixed, paraffin-embedded tissues were stained with hematoxylin and eosin (H&E, Carl Roth, Karlsruhe, Germany) and imaged on a Keyence Microscope BZ-X810 (Osaka, Japan). For immunofluorescence staining of tumor sections, samples were boiled for 30 min in 10 mM citric acid (AR6, PerkinElmer, Waltham, MA) to expose epitopes. Cell permeabilization was performed for 10 min with 0.25% Triton X-100 in PBS followed by blocking with 5% normal goat serum (Sigma-Aldrich) for 2 h at room temperature and overnight incubation with anti-CD3 (1:100, clone Rb19G5B7, Synaptic Systems, Göttingen, Germany) and anti-CD8 (1:100, clone GHH8/321E9, Synaptic Systems) at 4°C. After three washing steps with PBS, sections were incubated with anti-rabbit IgG H&L Alexa Fluor 488 (1:1000, Invitrogen) and anti-rat IgG H&L Alexa Fluor 555 (1:1000, Invitrogen) for 2h at room temperature. Upon washing with PBS, slides were mounted with ProLong Diamond Antifade Mountant with DAPI (Invitrogen). Images were acquired at ×200 magnification on a Nikon N-SIM confocal microscope (Nikon, Tokio, Japan) and then exported and processed in ImageJ (NIH, Bethesda, MD).

### Quantitative RT-PCR

Total RNA from splenocytes was isolated with NucleoSpin RNA kit (Macherey-Nagel). cDNA was synthesized using PrimeScript RT Reagent Kit (TaKaRa Bio Inc., Shiga, Japan). The qRT-PCR assays was performed using Luna Universal qPCR Master Mix (New England BioLabs, Frankfurt, Germany) on a CFX384 Touch Real-Time PCR Detection System (Bio-Rad Laboratories GmbH, Feldkirchen, Germany). *Ifi44* gene expression was normalized to expression of the housekeeping gene *Tbp1.* Primer sequences were as follows: *Ifi44*-F: 5’-GGCACATCTTAAAGGGCCACACTC-3’, *Ifi44*-R: 5’- CTGTCCTTCAGCAGTGGGTCATG-3’; *Tbp1*-F: 5’-TCTACCGTGAATCTTGGCTGTAAA-3’, *Tbp1*-R: 5’-TTCTCATGATGACTGCAGCAAA-3’).

### Western blot analysis

Splenocytes from *Trex1^Fl/Fl^ R26CreERT2 or Trex1^Fl/Wl^ R26CreERT2 mice were* isolated 14 days after TAM induction. Cells were lysed in RIPA buffer (50 mM Tris HCL, 150 mM NaCl, 1% SDS, 0.5% Na-deoxycholate, 1× Complete protease inhibitor (Roche)) for 30 min on ice. Lysates were centrifuged (16,000×g, 10 min, 4°C) and supernatant was incubated with Laemmli buffer at 95°C for 5 minutes. Proteins were separated on 12% SDS polyacrylamide gels and transferred to nitrocellulose membrane (Amersham Hybond-ECL, GE Healthcare). Membranes were blocked in 5% dried milk/TBS-T buffer and incubated over night with anti-mouse TREX1 antibody (clone C-11, Santa Cruz Biotechnology, Dallas, TX) and and anti-β-tubulin polyclonal antibody (Cell Signaling Technology, Danvers, MA). After washing with in Tris-buffered saline with Tween (TBS-T), membranes were incubated with HRP-conjugated rabbit-anti-mouse IgG (Cell Signaling Technology) for 1h at room temperature and finally washed again in TBS-T. For signal detection, membranes were incubated in Amersham^TM^ ECL^TM^ solution (Merck, Darmstadt, Germany) and images were captured on a Fusion FX - Western Blot & Chemi imaging instrument (VILBER, Eberhardzell, Germany).

## Acknowledgements

We thank Christa Haase and Sabine Ulbricht for expert technical assistance and Sarah Kaiser-Thom for help with animal husbandry. The study was supported by the German Research Foundation (DFG) through grant 369799452 – TRR237 Nucleic Acid Immunity, project B17 to A.R. and project B19 to R.B, and by funding from Bristol Myers Squibb. We thank the Dresden-concept Genome Center of the CMCB technology platform, TU Dresden, for excellent deep sequencing services.

