## Supplementary Figures for "Selective inflammation of the tumor microenvironment and invigorated T cell-mediated tumor control upon induced systemic inactivation of TREX1"

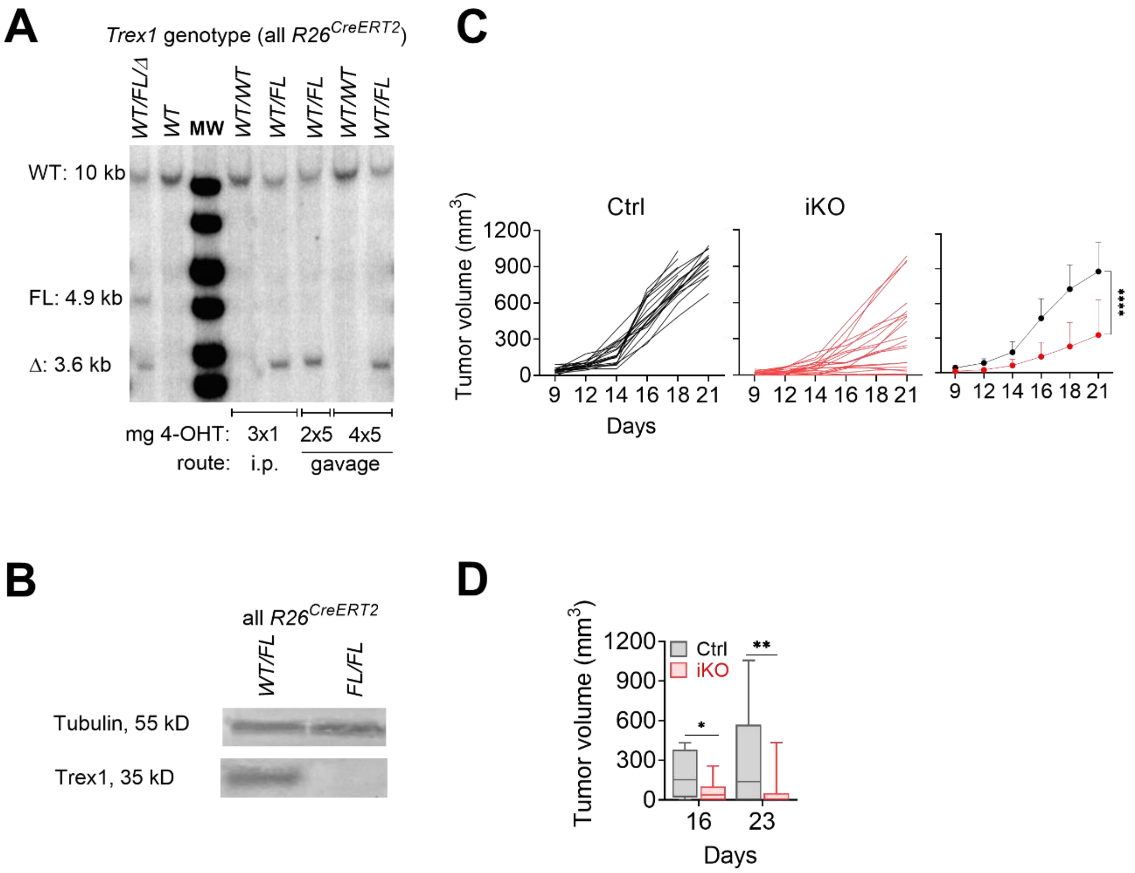


**Figure S1 (related to Fig. 1): Efficiency of induced *Trex1* deletion and growth of MB49 tumors in TREX1 iKO.**

**A)** Efficient deletion of the loxP-flanked *Trex1* allele induced by i.p. TAM injection (3× 1mg at daily intervals) or oral gavage (2× or 4× 5mg at daily intervals) in $Trex1^{FL/WT}R26^{CreERT2}$ mice. Southern blot analysis of DNA from total splenocytes 5 days after the last TAM administration. Splenocytes from $Trex1^{FL/WT}$ *CD19-Cre* mice served as a positive control containing WT, FL and deleted *Trex1* alleles. **B)** Western blot analysis of TREX1 expression in splenocytes 21 days after TAM induction (3× 1 mg at daily intervals), representative of 4 animals per genotype. **C)** Repetition of the experiment from Fig. 1 C and D. Monitoring of syngeneic B16.F10 tumor growth in male TREX1 iKO and control mice. Each line in the left two graphs represents one tumor. Data are summarized in the graph on the right. N=22, means ± SD, two-way ANOVA, ****p<0.0001. **D)** Tumor growth in female mice inoculated with $3\times10^5$ MB49 cells, N=15-24, pooled from two different experiments, means ± SD, two-way ANOVA, ****p<0.0001.

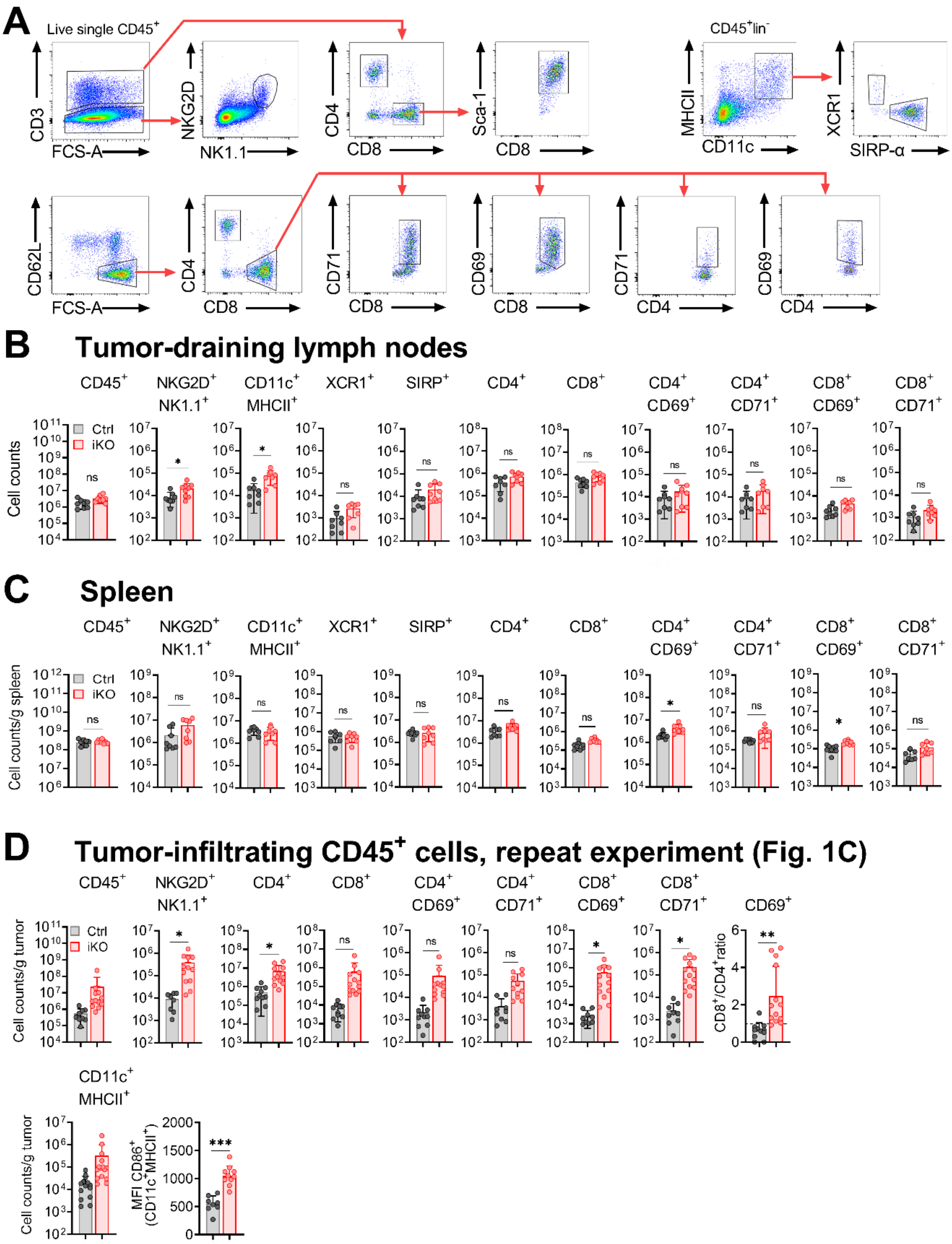


**Figure S2 (related to Fig. 2): Characterization of immune cell populations in tumor tissue, tumor-draining LN and spleen.**

**A)** Flow cytometric gating for quantification of individual immune cell populations. **B, C)** Flow cytometric determination of absolute numbers of immune cells per tumor-draining lymph node and absolute numbers of immune cells per gram of spleen of tumor-bearing mice sampled at day 21. Means ± SD, unpaired Student t test, ns – not significant, * $p<0.05$. **D)** Repetition of experiment in Fig. 1C. Flow cytometric determination of absolute numbers of immune cells per gram of tumor in single cell suspensions of tumors harvested on day 21. Means ± SD, unpaired Student t test, *$p<0.05$, **$p<0.01$, ***$p<0.01$.

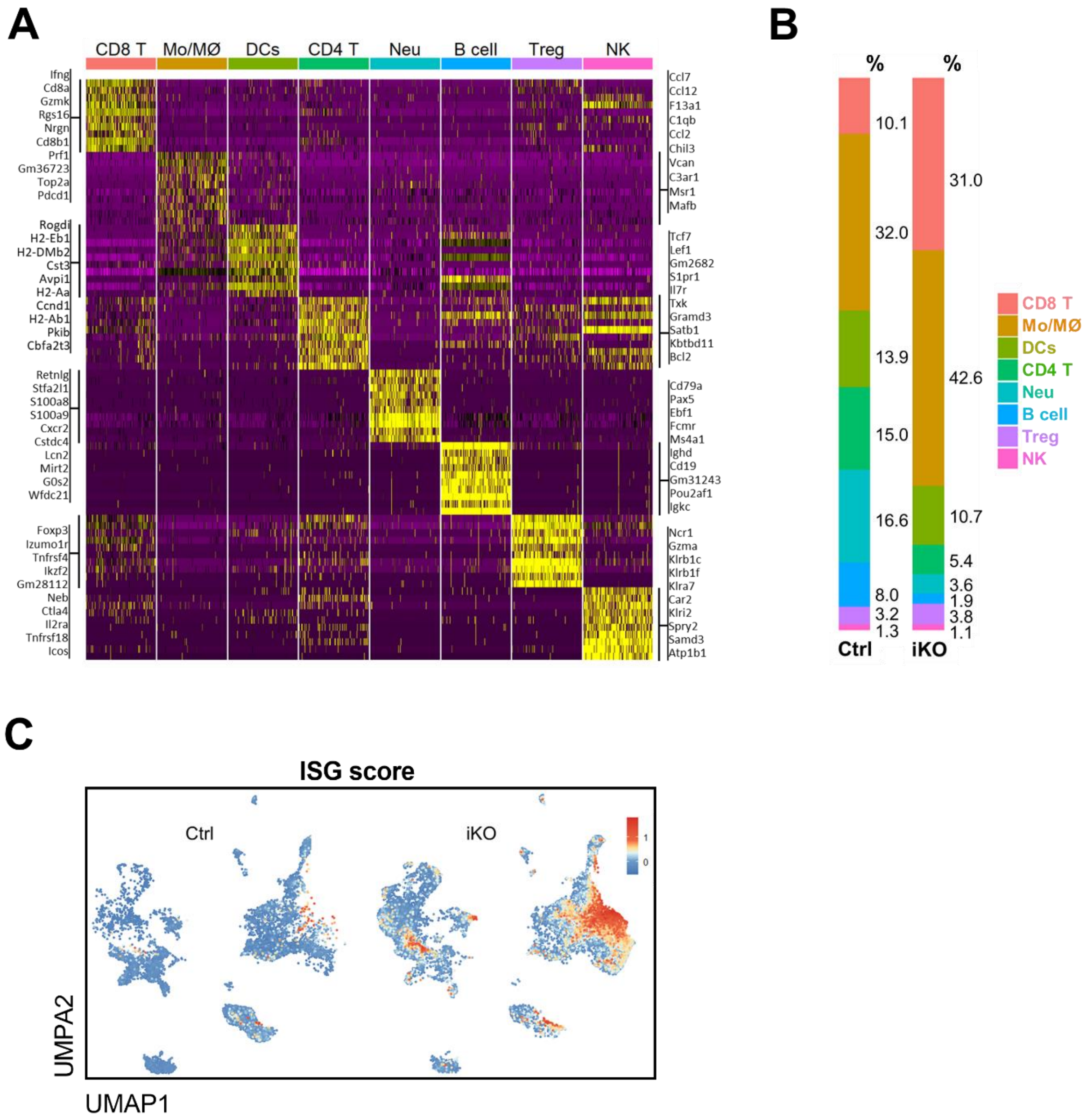


**Figure S3 (related to Fig.2): Single-cell RNA sequencing of CD45⁺ cells from tumors of TREX1 iKO and control hosts.**

**A)** Heatmap representing expression of genes defining UMAP clusters in the single-cell RNA-sequencing experiment in Fig. 2C. **B)** Frequencies of cells of the indicated clusters among total CD45$^{+}$ cells. **C)** Feature plot showing intensity of a transcriptional type I IFN-induced gene expression signature (score calculated from 40 ISGs, see Table S1)

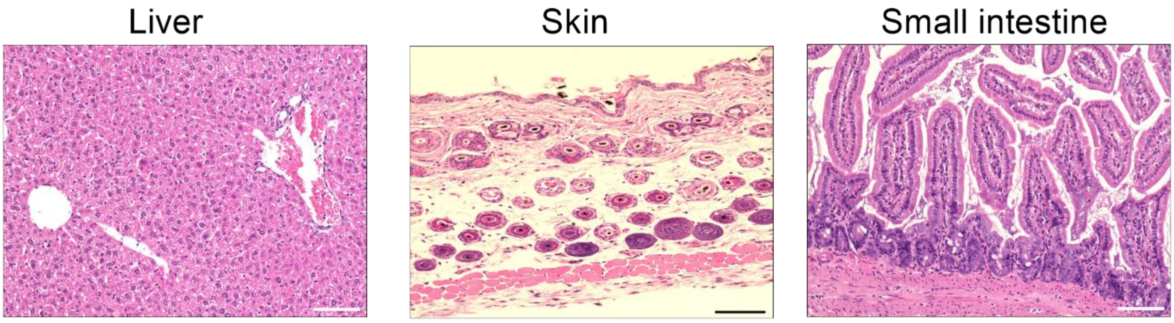


**Figure S4 (related to Fig. 2): No evidence for inflammation of non-tumor tissues of TREX1 iKO mice.**
H&E-stained sections of formalin-fixed, paraffin-embedded tissue sampled 5 weeks after TAM injection. Scale bars 100 µm.

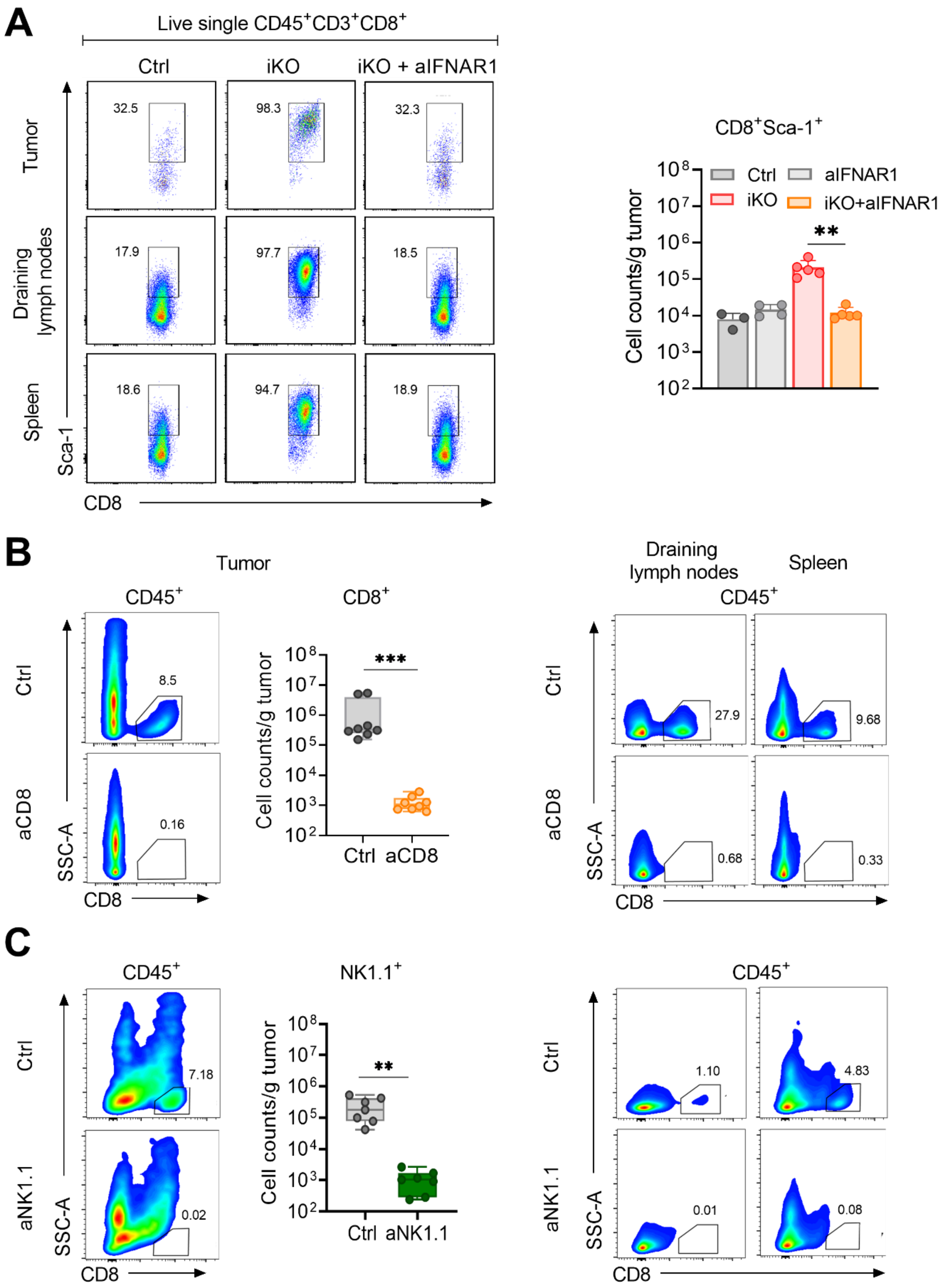


**Figure S5 (related to Fig. 3): Efficient IFNAR blockade and antibody-mediated depletion of CD8$^{+}$ T cells and NK cells.**

**A)** Expression of the IFN-induced surface protein SCA-1 on CD8$^{+}$ T cells and absolute numbers of SCA-1$^{+}$ CD8$^{+}$ T cells in control, TREX1 iKO or TREX1 iKO mice treated with anti-IFNAR1 antibody quantified by flow cytometry in tumor tissue, tumor-draining LN and spleen. Tissues were harvested on day 21 post tumor inoculation. A representative result (left) and summary of data (right) are shown. Means ± SD, One-way ANOVA followed by Sidak's multiple comparison test, ** $p<0.01$. Flow cytometric analysis of **B)** $CD8^+$ T cell and **C)** $NK1.1^+$ cell depletion in tumor tissue, in tumor-draining lymph nodes and spleen harvested on day 21 post tumor inoculation. Means ± SD, unpaired t test, ** $p<0.01$, *** $p<0.001$.

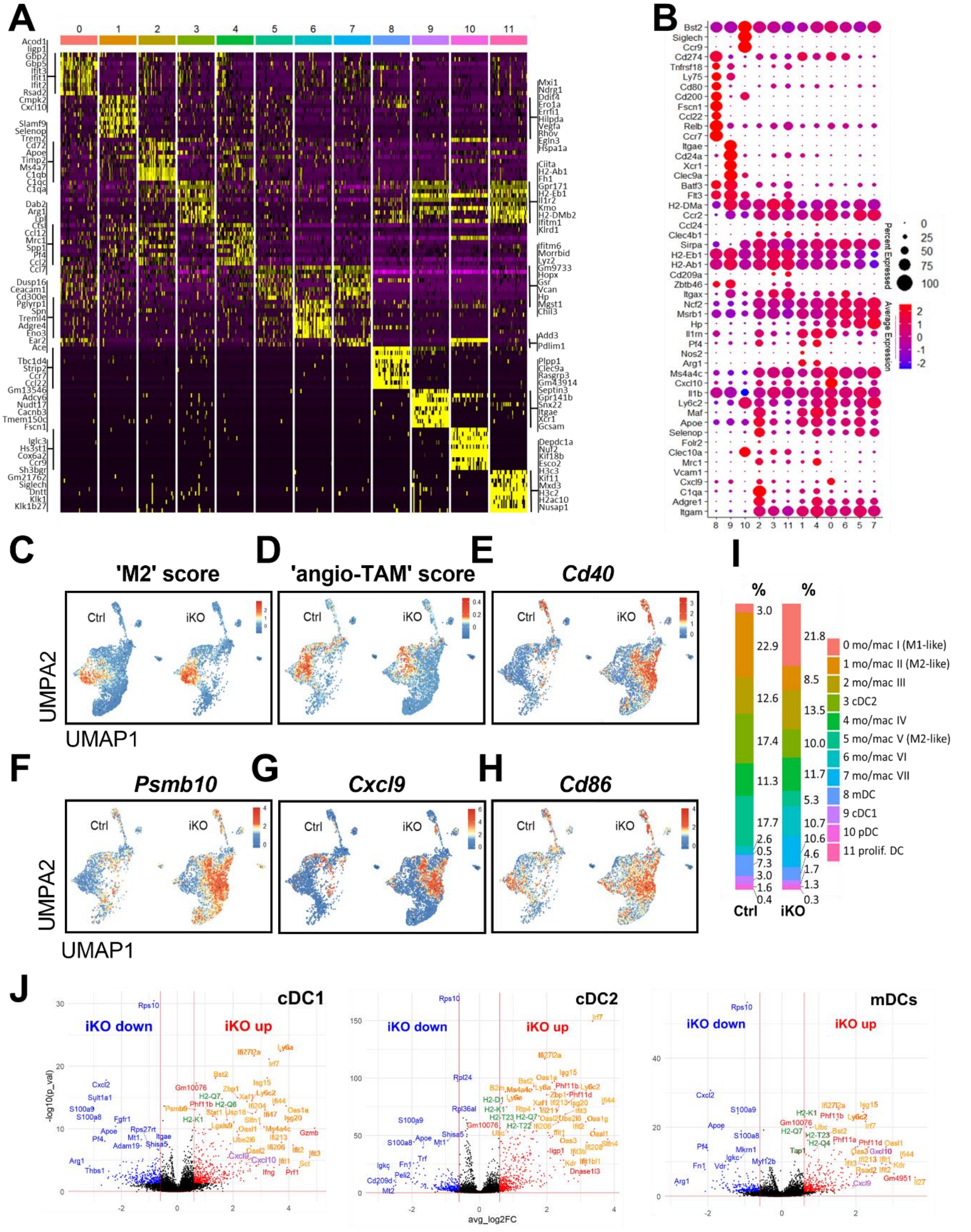


**Figure S6 (related to Fig. 4): Intra-tumoral myeloid cells switch toward more inflammatory phenotypes upon induced loss of TREX1.**

**A)** Gene expression heatmap of the single-cell RNA-sequencing data in Fig.4A showing expression of genes defining UMAP clusters. **B)** Expression of additional key marker genes. **C-H)** Feature plots showing the intensity of expression of **C)** M2-like macrophage signature (score was calculated from 98 genes, Table S1), **D)** ‘pro-angiogenic tumor-infiltrating macrophages’ signature as defined by Ma et al. (41) (score calculated from 5 genes, Table S1) **E)** *Cd40* **F)** *Psmb10*, **G)** *Cxcl9*, and **H**) *Cd86*. **I)** Frequencies of cell of the indicated clusters among the total sequenced myeloid cells. **J)** Volcano plot analysis of genes differentially regulated between cDC1, cDC2 and mDC populations from tumors of

TREX1 iKO versus control hosts. Names of ISGs indicated in orange, genes associated with antigen presentation in green. Note that cDC1s and mDCs strongly upregulate transcript levels of *Cxcl9* and *Cxcl10* encoding T cell recruiting chemokines (both also ISGs). All DC subsets downregulate the anti-inflammatory, type 2 associated enzyme arginase 1 (encoded by *Arg1*).

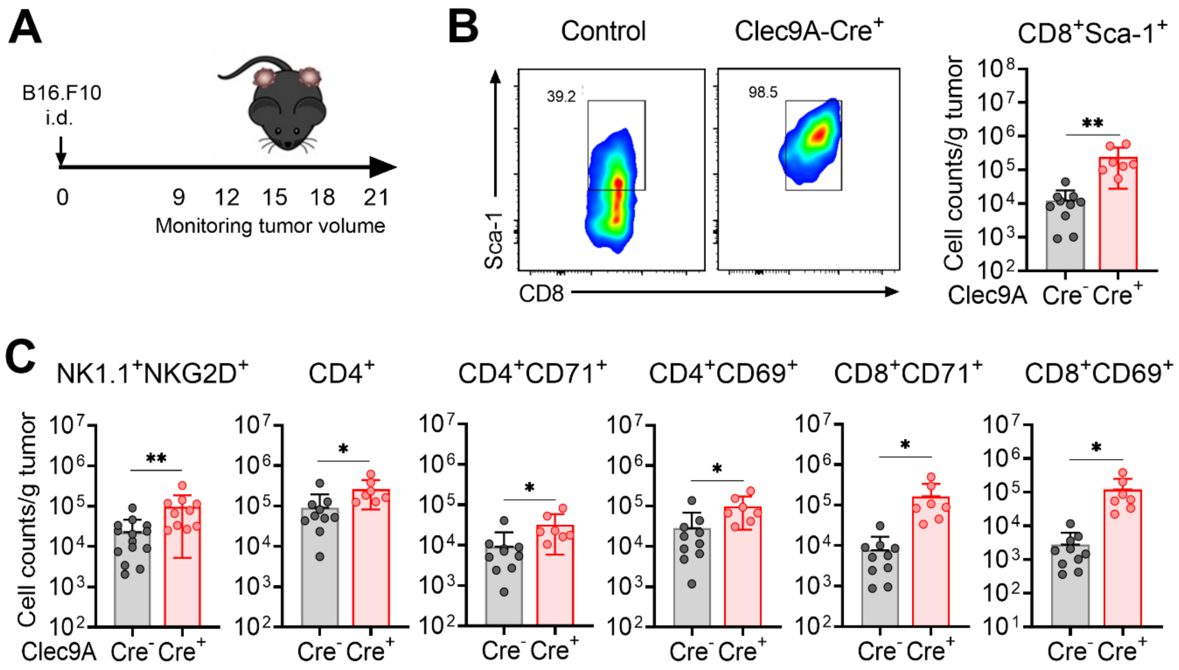


**Figure S7 (related to Fig. 4): Improved tumor control in hosts constitutively lacking TREX1 only in conventional dendritic cells.**
**A**) Experimental schedule of B16.F10 inoculation in *Trex1$^{FL/FL}$Clec9a-Cre$^{+}$* or *Trex1$^{FL/FL}$Cre-negative* control mice. **B)** Quantification of surface expression of SCA-1 encoded by the ISG *Ly6a* on CD8$^{+}$ T cells isolated from tumor tissue. Tumors were harvested on day 21 after tumor cell inoculation. Means ± SD, unpaired t test, ** p<0.01. **C)** Numbers of immune cells per gram of tumor tissue. Gating as in Fig. S2. Means ± SD, One-way ANOVA followed by Sidak's multiple comparison test, * p<0.05, ** p<0.01.

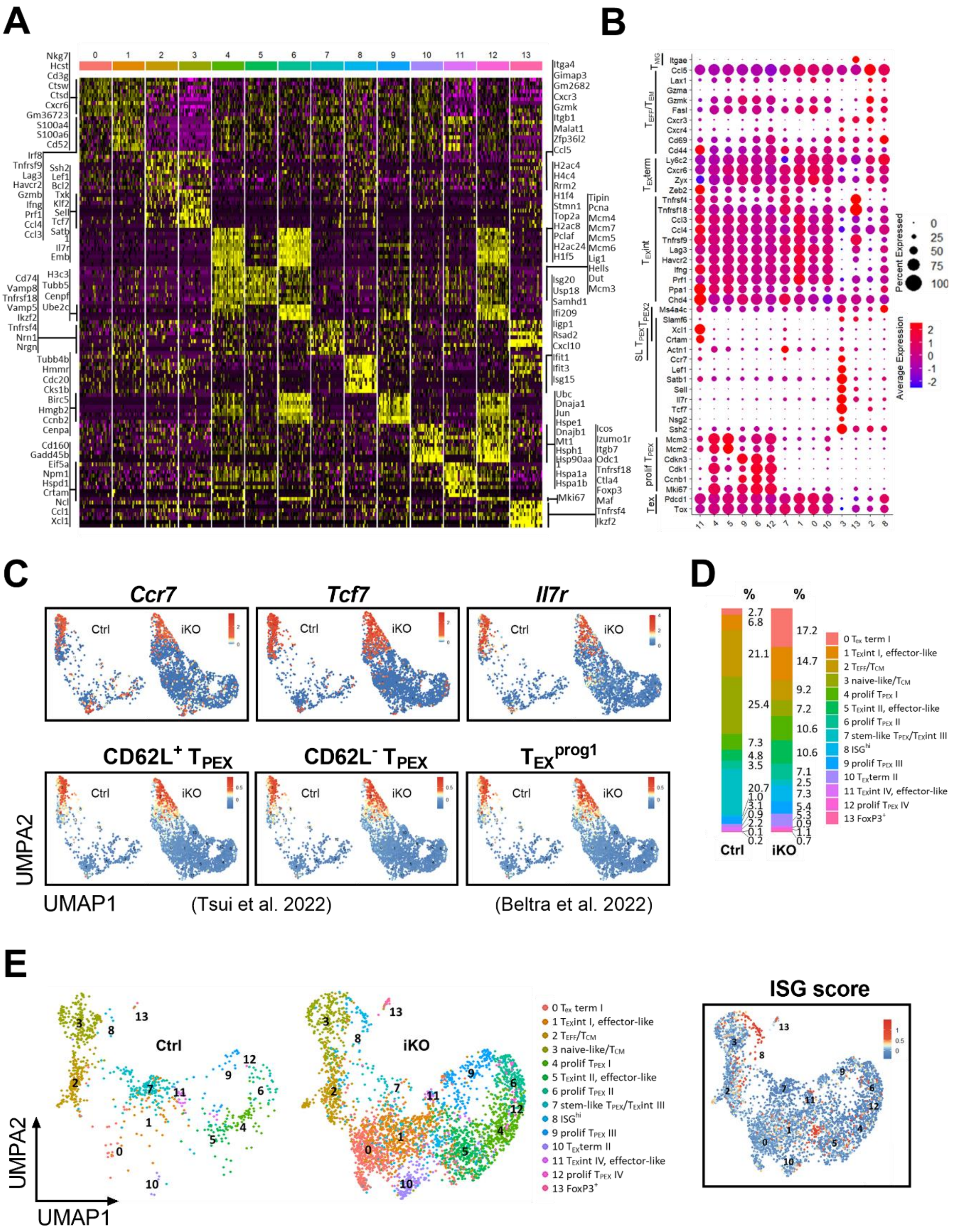


**Figure S8 (related to Fig. 5): Induced loss of TREX1 in host cells invigorates the intra-tumoral CD8$^+$ T cell response.**

**A)** Gene expression heatmap of the single-cell RNA-sequencing data in Fig. 5A showing expression of genes defining UMAP clusters. **B)** Dot plot of key marker genes. **C)** Feature plots showing expression of genes associated with stem-like $T_{EX}$ progenitors, *Ccr7*, *Tcf7* and *Il7r*, of genes defining the CD62L$^+$ and CD62L-negative populations of stem-like $T_{EX}$ progenitors (scores calculated from 72 and 26 genes, respectively, Table S1) described by Tsui et al. (45), and of genes defining $T_{EX}$ progenitor1 population as defined by Beltra et al. (46). **D)** Frequencies of cells of the indicated clusters among the total CD8$^+$ T cell population in tumors of control versus TREX1 iKO hosts. **E)** CD8$^+$ T cells (dataset of Fig. 5A) were re-clustered excluding the 40 top-expressed ISGs (Table S1). UMAP plots of re-clustered cells from control and iKO tumors (left) and feature plot showing the position of the $ISG^{hi}$ cells of Fig. 5A.

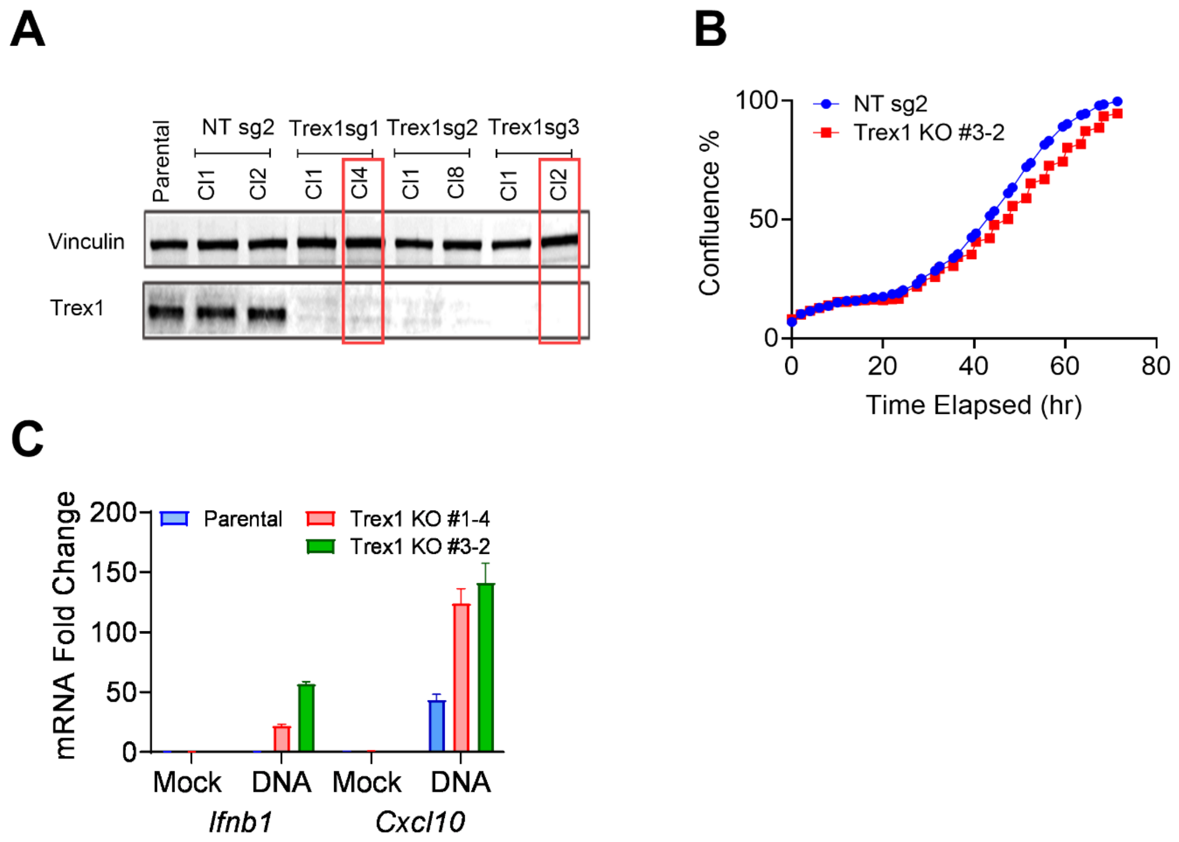


**Figure S9 (related to Fig.7): Loss of TREX1 in B16.F10 cells upon gene editing does not impair growth *in vitro* and does not trigger a spontaneous IFN response, but enhances responsiveness to transfected DNA.**

**A)** Upon RNP-based gene editing with one of three different guide RNAs and subcloning, parental cells and cells transfected with non-targeting (NT) guide RNAs along with gene-edited clones were analysed for TREX1 expression by western blotting. Editing of the Trex1 gene was also verified by sequencing (not shown). Clone #3-2 was used for all in vivo experiments. **B)** Kinetics of *in vitro* growth of control (NT) and the *Trex1*$^{-/-}$ clone #3-2 as determined by quantification of confluency through live cell imaging. **C)** IFN response of parental B16.F10 and *Trex1*$^{-/-}$ clones to transfection of calf thymus DNA. The transcript levels of *Ifnb1* and the ISG *Cxcl10* were determined by qRT-PCR. Fold change compared to mock transfected parental cells is shown.
